# Interactions of netrin-1 through its glycosylation sites immobilize Deleted in Colorectal Cancer (DCC) by favoring its constitutive clustering

**DOI:** 10.1101/2023.10.26.563740

**Authors:** Karen Uriot, Olivier Blanc, Nicolas Audugé, Orestis Faklaris, Olivier Blanc, Nathalie Chaverot, Evelyne Bloch-Gallego, Nicolas Borghi, Maïté Coppey-Moisan, Philippe P. Girard

## Abstract

Netrin-1 is a protein that attracts neurons expressing the membrane receptor Deleted in Colorectal Cancer DCC. In colon carcinoma, the interaction between netrin-1 and DCC prevents apoptosis. Crystallographic data suggest that these processes involve the clustering of DCC, the observation of which in cells remains elusive, as do the molecular determinants of DCC-netrin-1 interactions and their impact on DCC organization and mobility. To address these questions, we used fluorescence photobleaching, single-particle tracking and super-resolution techniques to characterize DCC organization and mobility on the cell surface. Our results show that netrin-1 impedes DCC mobility in the plasma membrane by promoting the growth of constitutive DCC nanoclusters at the expense of free DCC. Furthermore, we show that these effects are mediated primarily by the three glycosylation sites in the LamVI domain of netrin-1 and, to a lesser extent, by the C-terminal domain and its RGD binding site.

## Introduction

Netrins are secreted extracellular proteins that are involved in axon guidance (***Bloch-Gallego et al., 1999; Hedgecock et al., 1990; Serafini et al., 1996; Keino-Masu et al., 1996; Kolodziej et al., 1996; Fazeli et al., 1997; Tessier-Lavigne et al., 1988; Hamelin et al., 1993***). While initially identified for their ability to orient axons in the developing central nervous system, netrins are also involved in sculpting neuronal architectures and wiring the brain through the regulation of neuron branching (***Dent et al., 2004***), synaptogenesis (***Xu et al., 2004***) and axon regeneration (***Dun and Parkinson, 2017***). Moreover, netrins influence tissue morphogenesis and structural maintenance outside of the central nervous system by directing cell migration and regulating cell adhesion (***Hinck, 2004; Baker et al., 2006; Hebrok and Reichardt, 2004; Yebra et al., 2003; Marcos et al., 2009; Rappaz et al., 2016***), and are also implicated in tumorigenesis and cancer invasiveness (***Chen et al., 2017; Fitamant et al., 2008; Ylivinkka et al., 2016***). Netrin-1 is the prototype of the netrin family of laminin-related proteins. Its N-terminal sequence contains the Laminin VI domain and the Laminin V domain that consists in three epidermal growth factor (EGF) repeats (***Yamagishi et al., 2015***). Its positively charged C-terminal domain contains an RGD motif allowing binding to integrins (***Bányai and Patthy, 1999***). The receptor Deleted in Colorectal Cancer (DCC) is the generic receptor of netrin-1 involved in attractive responses in contexts of axon guidance, cell migration and tumorigenesis (***Fearon and Vogelstein, 1990***). DCC is a transmembrane protein characterized by the presence of four Ig domains and six fibronectin type III (FNIII) repeats in its extracellular domains (***Hedrick et al., 1994***). Additionally, DCC intracellular domain contains a caspase-3 proteolysis site at Asp 1290 (***Mehlen et al., 1998***). Functional studies in C. elegans have shown that both the laminin VI, EGF-2 and EGF-3 repeat domains of netrin-1 are required for dorsal and ventral axon migrations (***Lim and Wadsworth, 2002***). Structural studies have revealed three sites of interaction between Netrin-1 and DCC in the crystalized complex of netrin-1 deleted of its Cterminal domain and different isolated fragments of DCC (***Xu et al., 2014; Finci et al., 2014***). However, only some of these interactions were confirmed in biochemical assays (***Geisbrecht et al., 2003***). Nevertheless, a model based on netrin-1 interactions with FNIII repeats of DCC proposes that netrin-1 signaling emerges from netrin-1-induced DCC homodimerization followed by large-scale clustering (***Finci et al., 2014, 2015***). On the other hand, fluorescence microscopy studies suggest that exposure to netrin-1 increases cell surface levels of DCC (***Gopal et al., 2016***) and micrometer-scale clustering at the surface of cortical axons (***Bouchard et al., 2004; Matsumoto and Nagashima, 2010***). However, the roles of the different netrin-1 domains in the interaction with full length DCC in a physiological cell context remain to be determined.

Here, we address this question by investigating the effects of various mutants of netrin-1 on the mobility and the nanoscale organization of DCC in human HEK 293T embryonic kidney cells and in human Caco-2 colorectal cancer cells, two model cell lines of netrin-1/DCC signaling that do not endogenously express netrin-1 or DCC (***Mehlen et al., 1998***). Altogether, our results converge toward a model by which DCC mobility emerges from its constitutive nanoscale clustering, which is promoted by netrin-1 interaction primarily through its glycosylation sites and to a lesser extent through its RGD site.

## Results

### Recombinant netrin-1-mCherry is chemoattractive and captured by neurons

To visualize netrin-1 in live cells, we made a recombinant full-length netrin-1 tagged with mCherry at its C-terminus (FL, Fig. 1A). We measured its attracting effect on neurons in a chemotaxis assay between mouse rhombic lip explants and aggregates of either HEK EBNA cells stably expressing unlabeled recombinant netrin-1 (HEK NTN1, Fig. 1B and C), or HEK293T stably expressing full-length netrin-1 tagged with mCherry (HEK FL, Fig. 1D). As expected, the surface area covered by axons elongating from spinal cord explants as well as the surface area occupied by nuclei performing nucleokinesis were both larger towards the HEK NTN1 aggregate (proximal growth and nucleokinesis) than in the opposite direction (distal growth and nucleokinesis) (Fig. 1E-F). Moreover, we found that the neural explant exhibited the same behavior toward HEK FL cells. This confirms that FL is functional at attracting neurons. Moreover, we found that FL colocalized with proximal but not distal axons from the neural explants (Fig. 1D). This shows that neurons capture FL secreted by HEK cells.

**Figure 1.**
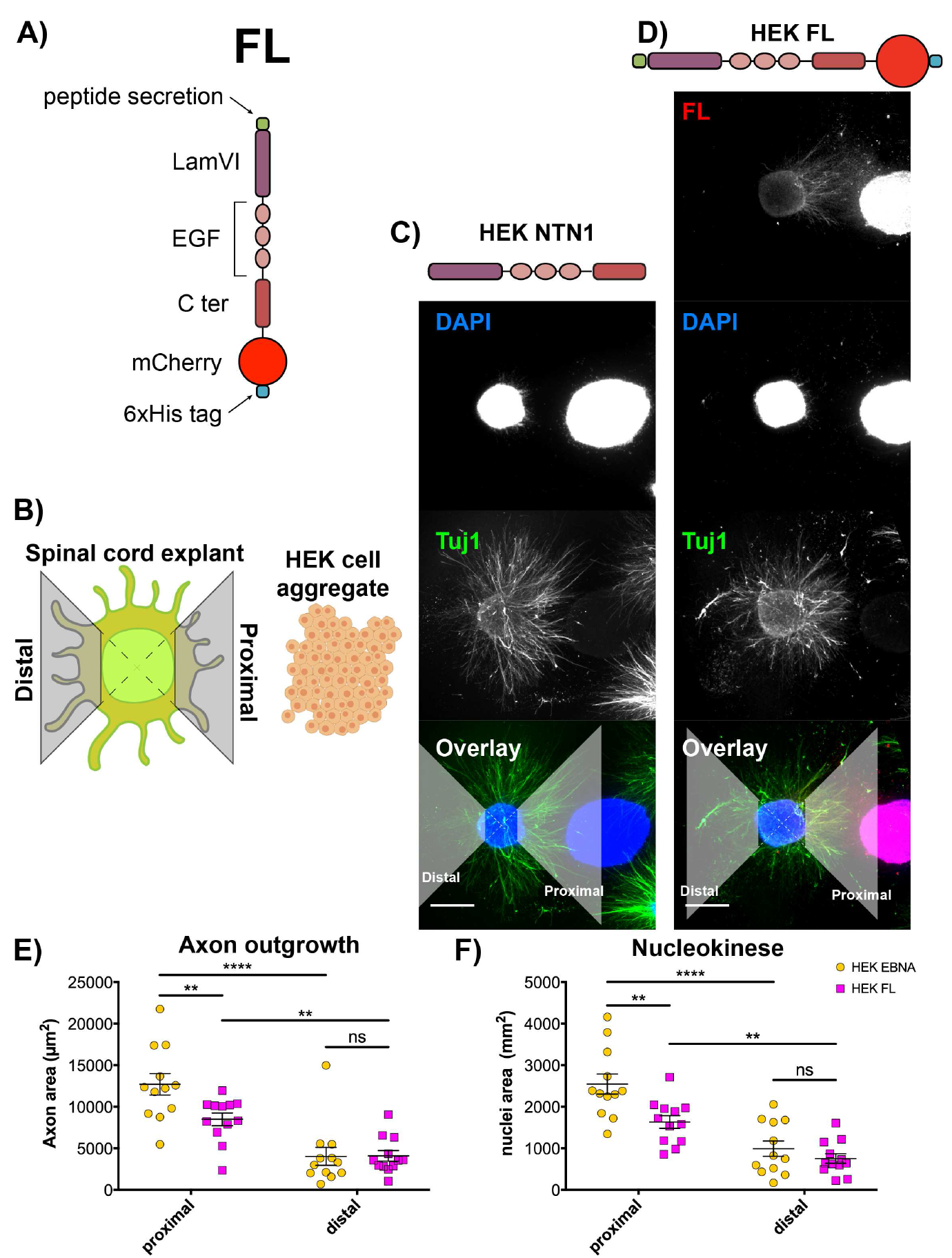
(A) Full length netrin-1 construct (FL). LamVI = Laminin G domain, EGF = epidermal growth factor (EGF)-like repeat, Cter = C-terminal domain, 6xHis tag = six consecutive histidine residues. (B) Principle of quantification of neuritic outgrowth and neuronal migration. Proximal and distal quadrants (grey) for neuritic processes and cell bodies quantification. (C-D) Rhombic lip explants from E12 mouse embryos cocultured 72 hours with aggregates of HEK EBNA cells secreting unlabeled recombinant netrin-1 (HEK NTN1) (C) or HEK cells secreting full-length netrin-1 tagged with mCherry (HEK FL) (D). Scale bar: 500 μm. (E) Surface area covered by neuritic processes in the proximal and distal quadrants. (F) Surface area covered by cell nuclei in proximal and distal quadrants. Mean ± SEM, ns *P* > 0.05, **** *P* < 0.0001, ** *P* ≤ 0.01, Student’s paired t-tests.

### Netrin-1 glycosylation is the major determinant of its recruitment by DCC at the cell surface

To address the roles of netrin-1 domains and post-translational modifications on its capture by its receptor DCC, we derived deleted or mutated variants from the FL construct stably expressed in HEK cells: a variant deleted of its Lamin VI domain (dLamVI) and one where we mutated the four putative N-glycosylated asparagines (AAC) to alanines (GCC) in the LamVI (N97, N118 and N133) and LamV-3 (N419) domains (glycoM), one deleted of its C-terminus (dC) and one with an RGE sequence replacing the integrin-binding RGD in the C-terminus (RGE) (Fig. 2A).

**Figure 2.**
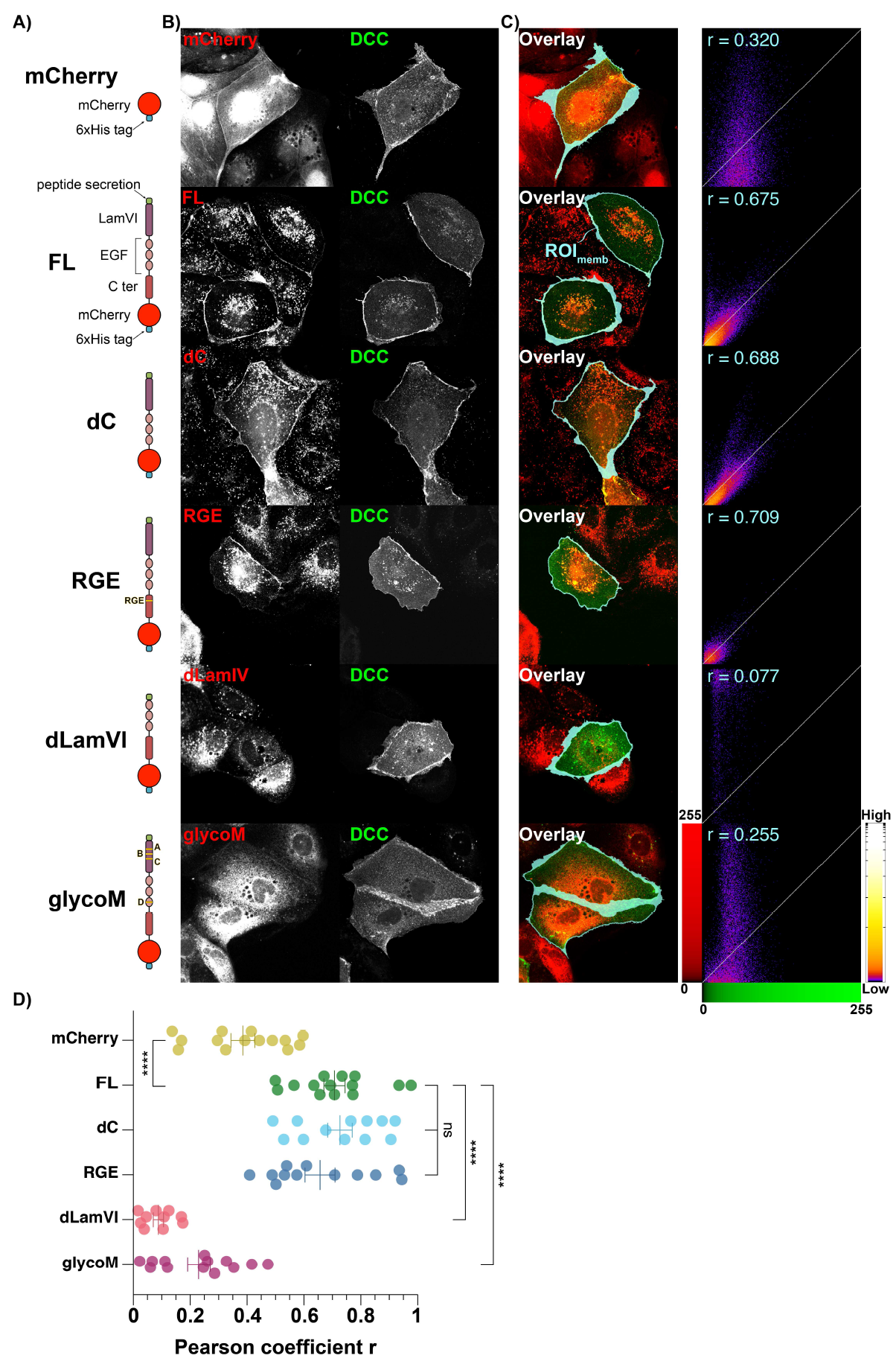
(A) Constructs used in this study: mCherry, and FL and its mutants (dC, dLamVI, glycoM and RGE) fused to mCherry. (B) Representative confocal images of Caco-2 cells stably expressing constructs in (A) transiently transfected with DCC-GFP. Scale bar: 20 μm. (C) Overlay of (B) with ROI_memb_ (cyan, see Materials & Methods) and its pixel fluorogram, r: Pearson’s correlation coefficient. (D) Pearson’s coefficients r (11 < *n* < 15 cells per construct, mean ± SE), **** *P* < 0.0001. Unpaired two-tailed Mann–Whitney test.

To validate the secretion of netrin-1 variants and their capture by cells expressing DCC, we quantified by Fluorescence Activated Cell Sorting (FACS) the proportion of “green”-positive and “red”-positive cells in co-cultures of HEK cells expressing them or mCherry alone together with HEK cells expressing GFP alone or DCC-GFP (Fig. S1, see Materials and Methods). We observed the capture of all netrin-1 variants by cells expressing DCC-GFP and to a larger extent than by cells expressing just GFP (Fig. S1C). Futhermore, in co-cultures with FL, dC and RGE variants, the amount of netrin-1 variant recruited by DCC-GFP cells more strongly correlated with the expression level of DCC-GFP than it did in co-cultures with mCherry alone, dLamVI or glycoM. It is worth noting that the amount of any netrin-1 variant recruited by GFP cells was low and did not correlate with GFP levels. These findings support that a DCC-specific capture of netrin-1 occurs and that it requires intact putative glycosylation sites of netrin-1, while its C-terminus and the RGD peptide are dispensable.

Then, we examined the relative subcellular localizations of netrin-1 and DCC by confocal microscopy in fixed Caco-2 cell lines expressing netrin-1-mCherry variants or mCherry alone and transiently transfected with DCC-GFP, to mimic autocrine signaling. We found that DCC-GFP localized mainly at the plasma membrane and to some extent in intracellular vesicles, in all conditions, even in the absence of netrin-1 (Fig. 2B). This shows that the presence of DCC at the plasma membrane does not require cell exposure to netrin-1. Moreover, we found that at the plasma membrane, the levels of FL, dC and RGE variants scaled with that of DCC-GFP, while it was not the case for mCherry, dLamVI or glycoM (Fig. 2C). In agreement with FACS of co-cultures, this supports that an interaction with DCC and netrin-1 occurs at the plasma membrane and that it requires intact putative glycosylation sites of netrin-1, while its C-terminus and the RGD peptide are dispensable.

### Netrin-1 impairs DCC mobility at the minute time-scale by interacting with DCC

To assess the impact of netrin-1 on DCC mobility at the minute time-scale, we conducted fluorescence recovery after photobleaching (FRAP) measurements on DCC-GFP (Fig. 3A) in Caco-2 cells co-expressing the FL construct or the mCherry construct as a control. Control experiments were also performed on DCC-GFP cells in presence of the AF5 antibody known to prevent the interaction between netrin-1 and DCC (***Keino-Masu et al., 1996***), or on cell expressing DCC-GFP D1290N which cleavage by caspases is prevented (***Mehlen et al., 1998***) (Fig. 3B). For all conditions, DCC-GFP fluorescence recovered to a stable plateau within minutes (Fig. 3C). A fit with a reaction-diffusion model showed that FL increased the immobile fraction of DCC-GFP and to a lesser extent the reaction fraction at the expense of the diffusive fraction (Fig. 3C-D). Addition of the AF5 antibody reverted the immobile and diffusive fractions of DCC-GFP in cells exposed to netrin-1 close to that in the absence of netrin-1 (Fig. 3C-D), as expected. Overall, this shows that netrin-1 substantially stabilizes DCC at the plasma membrane by its interaction with DCC.

**Figure 3.**
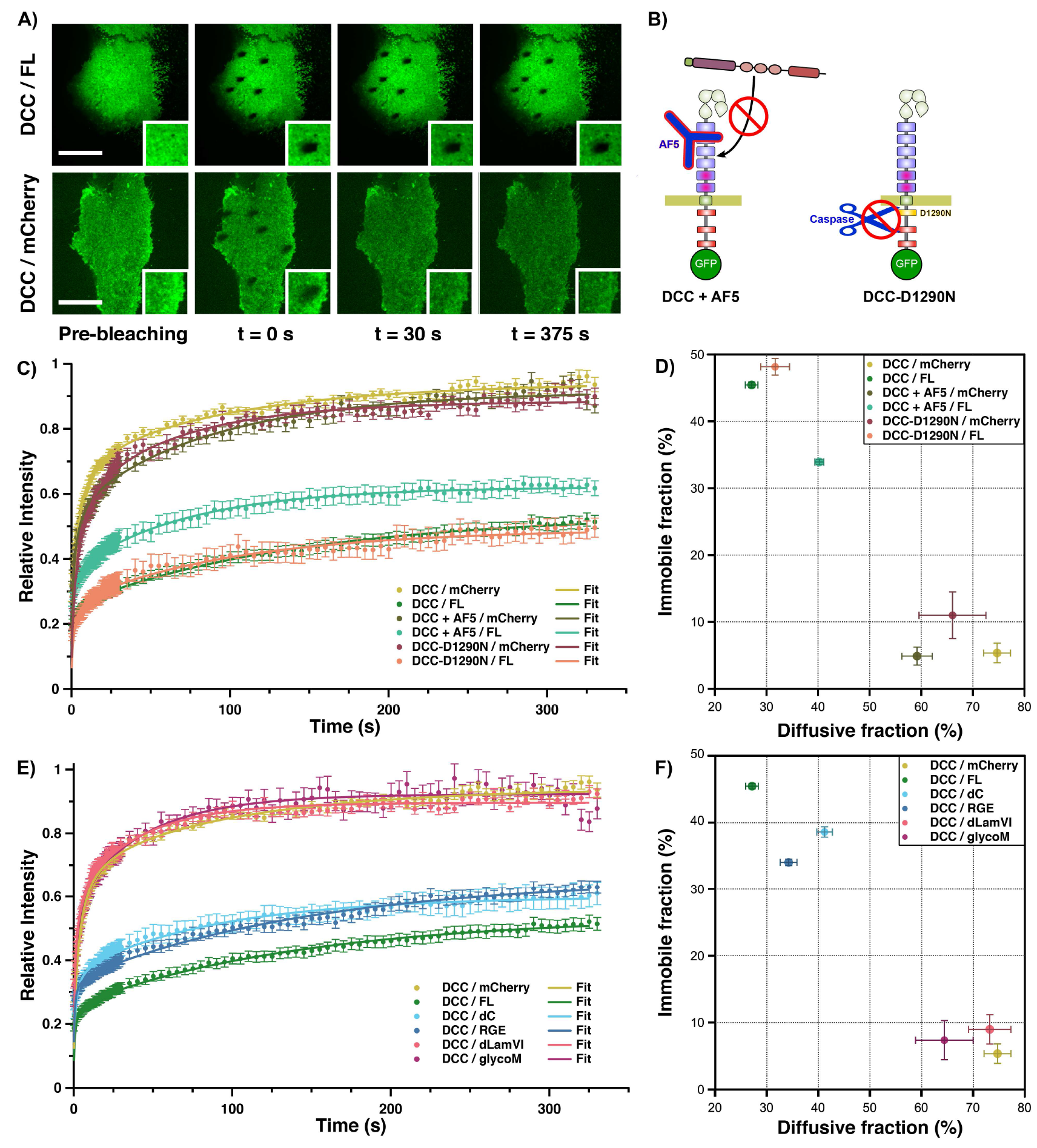
(A) Representative images of DCC-GFP in Caco-2 cells expressing either FL (top) or mCherry (bottom) construct during a FRAP experiment. Scale bar: 10 μm. (B) Left: AF5 antibody inhibiting the netrin-1 binding site of DCC. Right, DCC-D1290N inhibiting cleavage by caspases. (C) Fluorescence recovery curves of DCC-GFP or DCC-D1290N-GFP in Caco-2 expressing netrin-1 FL or mCherry with or without AF5 antibody (*n* > 18 cells per condition, mean ± SEM) (D) Immobile fractions vs. diffusive fractions from (C). (E) Average fluorescence recovery curves of DCC-GFP in Caco-2 cells expressing FL, dC, dLamVI, glycoM, or mCherry (*n* ≥ 18 cells per condition, mean ± SEM). (F) Immobile fractions vs. diffusive fractions from (E).

In the absence of netrin-1, DCC can be cleaved by caspases (***Forcet et al., 2001***). In that case, the GFP-tagged cytoplasmic tail of DCC would substantially contribute to its diffusive fraction. However, we found that the D1290N mutation only marginally affected the immobile and diffusive fractions of DCC-GFP with or without netrin-1 (Fig. 3C-D). This shows that most of DCC is constitutively full length, regardless of netrin-1 presence, and that DCC immobilization by netrin-1 does not result from cleavage prevention but the trapping of the whole receptor.

Netrin glycosylation is the major determinant of DCC mobility impairment by netrin Next, we assessed the roles of netrin-1 domains on DCC mobility. First, we found that the presence of the dLamVI and glycoM mutants led to DCC-GFP mobility resembling that seen without FL. Specifically, the immobile fraction decreased significantly, while the diffusive fraction increased in comparison to DCC-GFP with FL (Fig. 3E-F). These results highlight the essential role of the putative glycosylation sites within netrin-1 for immobilizing DCC, particularly those in the Laminin VI domain. Comparatively, the decrease in DCC-GFP immobile fraction and the increase in diffusion fraction were less pronounced when exposed to the dC and RGE mutants instead of FL (Fig. 3E-F). These outcomes signify that the C-terminus of netrin-1 and the RGD sequence contribute to a significant yet moderate extent to DCC immobilization.

### Minute time-scale DCC mobility mostly arises from that at the second time-scale

To better resolve the scale at which DCC immobilization by netrin-1 occurs, we performed single particle tracking (SPT) of DCC-GFP exposed to netrin-1 and its mutants (see Materials & Methods). In all conditions, DCC-GFP exhibited an immobile fraction of receptors that remained within a surface of about 20 × 10^3^ nm^2^ for at least 500 ms, a confined fraction that remained within a surface of about 80 × 10^3^ nm^2^ within the same time frame, and a free population that diffused without spatial limits (Figs 4A-B and S2). We observed that the diffusion coefficient of free DCC lied around 0.06 μm/s in the absence of netrin-1 or in the presence of the dLamVI or glycoM mutants, while it dropped to about twice less in the presence of FL or the dC and RGE mutants (Fig. 4C). Additionally, the confined fraction of DCC remained within a surface of about 80 × 10^3^ μm^2^ in the absence of netrin-1 or in the presence of the dLamVI or glycoM mutants while it dropped below 40 × 10^3^ μm^2^ in the presence of FL or the dC and RGE mutants (Fig. 4D).

**Figure 4.**
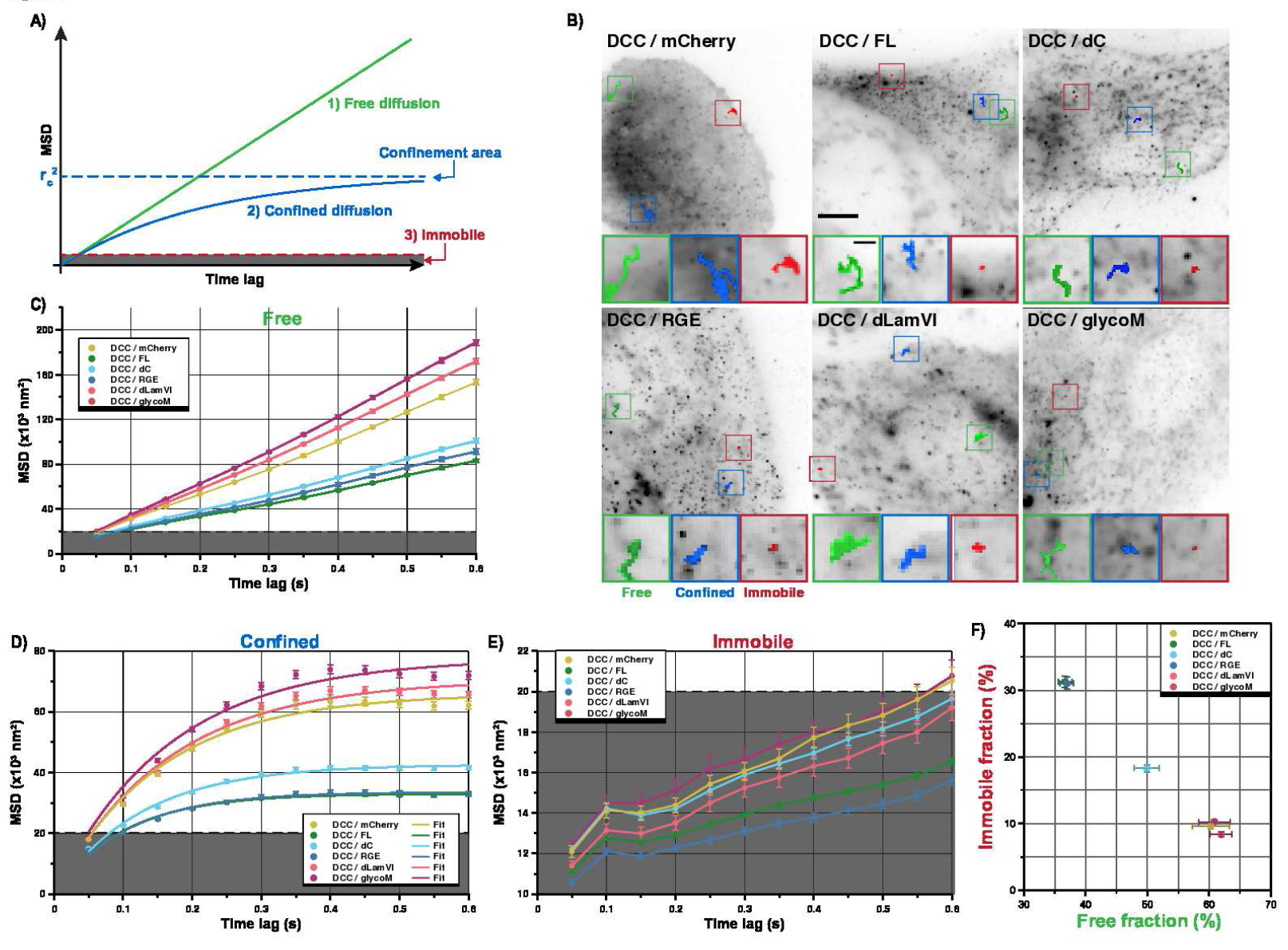
(A) Classification of single-molecule trajectories into free diffusion (green), confined diffusion (blue) and immobile (red) based on their MSD. (B) Representative images of DCC-GFP in Caco-2 cells expressing FL, dC, RGE, dLamVI, glycoM or mCherry with typical tracks of a particle for each diffusion mode (same colors as in A). Scale bar: 5 μm (1 μm in zoomed-in panels). (C-E) MSD vs. time lag of free (C), confined (D) and immobile (E) particles of DCC-GFP in cells above (5000 tracks from *n* > 10 cells per condition). (F) Immobile fractions vs. free fractions from the same data as (C-E). Mean ± SEM.

Consistently, in the absence of netrin-1, about 10% of DCC was immobile, 60% free and the rest confined, while in the presence of FL, the immobile fraction raised to more than 30% and the free fraction dropped to less than 35% (Fig. 4F). In the presence of the dLamVI or glycoM mutant instead, DCC mobility reverted to that in cells not exposed to netrin-1 (Fig. 4F). Conversely, the dC mutant had a milder effect, with immobile and free fractions of DCC falling around 18 and 50%, respectively, while the effect of the RGE mutant was not substantially different from that of FL (Fig. 4F).

Overall, the SPT results follow a similar trend as the FRAP results for most conditions (Fig. S3): DCC immobilization mostly relies on the putative glycosylation sites in the laminin VI domain of netrin-1, while the C-terminus plays a more moderate role. The RGD sequence, however, does not appear to affect the second scale mobility of DCC (Fig. 4), while it does affect its minute scale mobility (Fig. 3). This indicates that the mechanisms underlying the control of DCC mobility by netrin-1 occur at the second scale and micrometer scale, except for those that involve the RGD sequence, which occurs at larger scales.

### Netrin-1 increases constitutive nano-clustering of DCC through its glycosylation sites and, to a lesser extend it RGD site

To investigate whether netrin-1 influences DCC mobility through changes in DCC clustering within the plasma membrane, we imaged single-molecule localizations of DCC-PAGFP by Photo-Activated Localization Microscopy (PALM) in fixed cells stably expressing netrin-1 variants (see Materials & Methods) (Fig. 5). We first observed that regardless of netrin-1 exposure, DCC formed clusters, and the number of these clusters increased proportionally with DCC surface density (Fig. 5E-F-G). This finding underscores that DCC nano-clustering is constitutive and concentration-dependent. Moreover, we found that cluster cardinality, surface area and the fraction of DCC involved in clusters increased with DCC surface density, although in a netrin-1-dependent manner (Fig. 5). Specifically, the presence of FL led to the steepest increase, followed by the dC and RGE mutants. In contrast, the dLam and glycoM mutants exhibited a moderate increase akin to that seen in the absence of netrin-1 (Fig. 5E-F-G). These findings indicate that even though cluster number is independent of netrin-1 (Fig. 5D), netrin-1 favors bigger and larger clusters at the expense of free DCC. This effect is primarily driven by the glycosylation sites within netrin-1 and, to a lesser extent, by the RGD site in its C-terminal domain.

**Figure 5.**
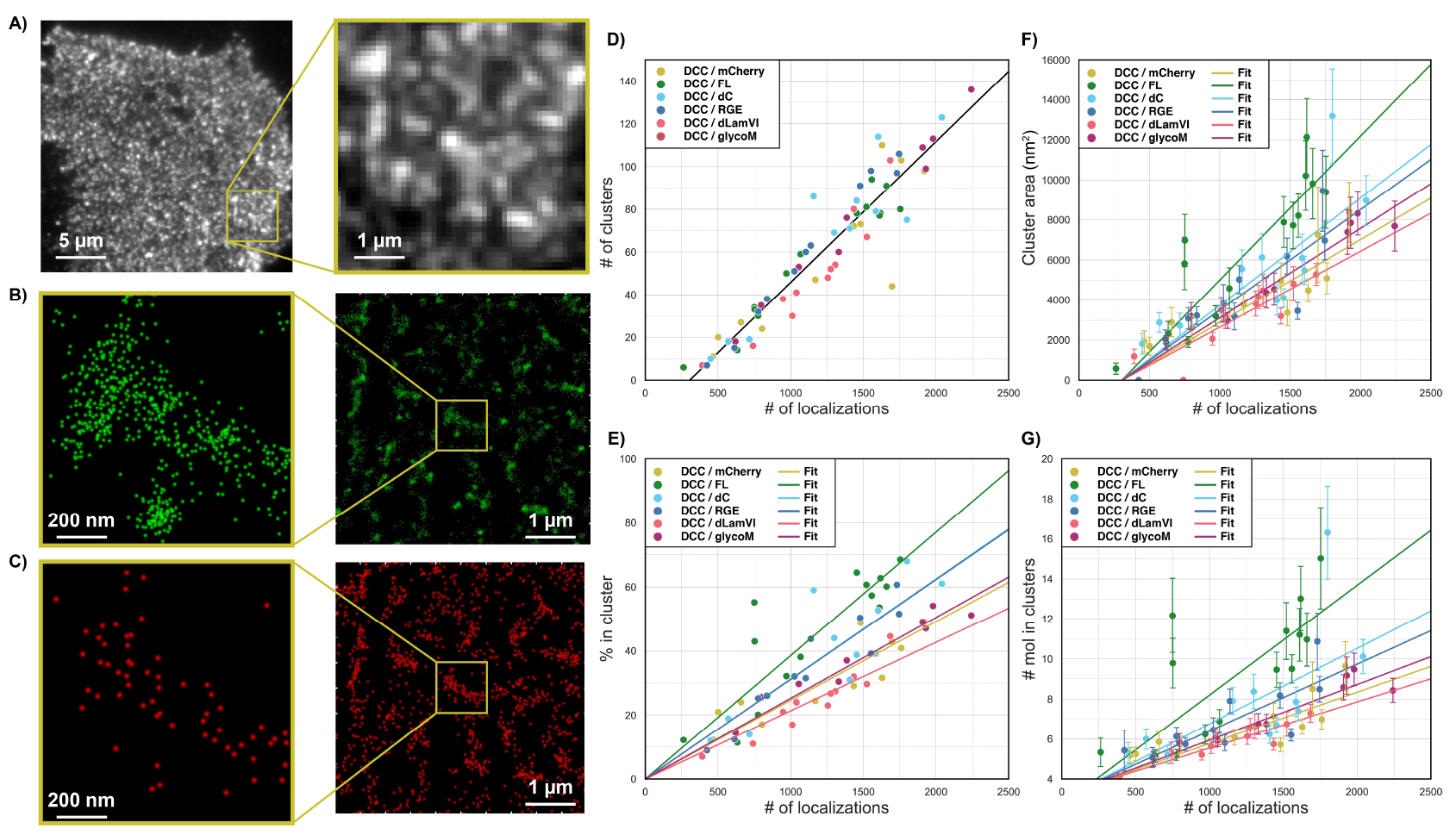
(A) Cumulative intensity of 2000 PALM frames of DCC-PAGFP in HEK FL cells. (B) PALM localizations of DCC-PAGFP before blinking correction from the acquisition shown in (A). (C) Single DCC-PAGFP molecule localizations within the same area after blinking correction. (D-G) DBSCAN analysis of (D) cluster number vs. localization number, (E) fraction of localizations in cluster vs. localization number, (F) cluster area vs. localization number, (G) cluster cardinality vs. localization number. Mean ± SEM (*n*_FL_ = 14, *n*_dC_ = *n*_RGE_ = *n*_mCherry_ = *n*_dLamIV_ = 11, *n*_glycoM_ = 9 cells).

Overall, the impact of netrin-1 on nano-scale clustering of DCC exhibited a strong correlation with its influence on DCC mobility at the second and minute time-scales (Fig. S4). Consequently, it is possible to discriminate the two dLam and glycoM mutants, which display pronounced effects abolishing most of netrin-1 properties, and the RGE and dC mutants, which induce comparatively modest alterations (Fig. S4C).

## Discussion

Our results show that DCC clusters are constitutive and concentration-dependent (Fig. 5D). Therefore, netrin-1 is dispensable for DCC clusters nucleation and maintenance. Consequently, DCC clusters must rely on interactions that do not involve netrin-1. For a given concentration of DCC, addition of netrin-1 results in an increase in the fraction of DCC receptors in clusters and consequently an increase in cluster cardinality and area (Fig. 5E-G), without increase in cluster number (Fig. 5D). Thus, netrin-1 promotes the growth of existing clusters rather than their fusion, or the nucleation of new ones. Cluster size depends on receptor concentration but never exceeds about 20 × 10^3^ nm^2^ in the presence of netrin-1 and twice less without (Fig. 5F). In SPT experiments, this is the range within which immobile particles stay below the second time-scale, while particles that experience confined diffusion explore larger areas that decrease, rather than increase, upon addition of netrin-1 (Fig. 4D). These results are thus consistent with a model by which clusters contain immobilized DCC receptors and confine mobile receptors travelling nearby in between them. Consistently, the fraction of immobile receptors grows upon addition of netrin-1 in both FRAP and SPT experiments while the confined fraction of receptors in SPT and fraction of receptors that experience reaction in FRAP remain mostly unaffected upon addition of netrin-1 (Fig. 3F). Since netrin-1 addition neither affect the number of clusters, we speculate that the fraction of confined particles is in fact the population of receptors that experience transient interactions from a cluster to another and that will scale in proportion to the number of clusters. Interestingly, since the fraction of confined receptors and the number of clusters remain unchanged while netrin-1 decreases the area of confinement, we predict that the surface density of confined receptors must increase. Ultimately, clusters grow upon addition of netrin-1 at the expense of the population of freely diffusing particles, as if netrin-1 favored the immobilization of receptors previously in transient interaction that previously free receptors would replace. We summarize this model in Fig 6. While our results showing that netrin-1 favors DCC clustering are consistent with crystallographic data (***Xu et al., 2014; Finci et al., 2014***), the existence of constitutive clusters in the absence of netrin-1 reveals other direct or indirect interactions that are instrumental for receptor clustering. This may be due to DCC residing in lipid rafts (***Guirland et al., 2004***), as other axonal guidance receptors, since lipid raft localization of DCC was shown to be required for netrin-1-induced DCC signaling (***Hérincs et al., 2005; Petrie et al., 2009***).

**Figure 6.**
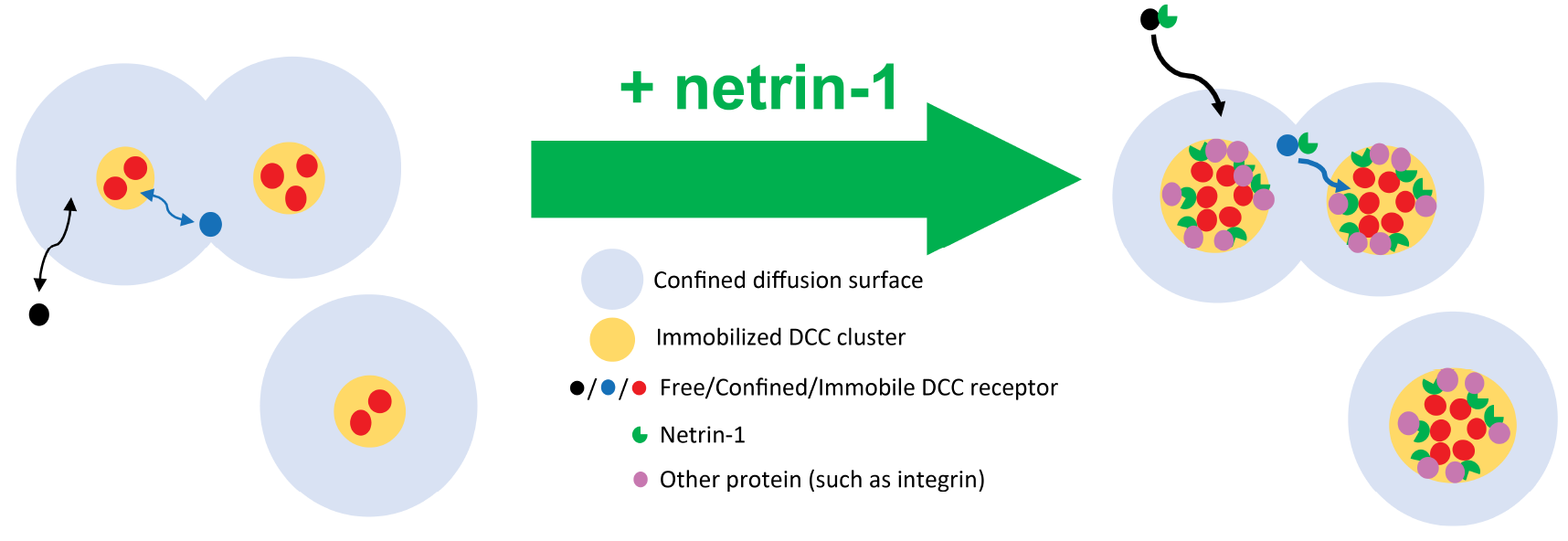
Working model of the effect of Netrin-1 on DCC spatial organization and mobility. The DCC receptors display either free diffusing (black), confined diffusing (blue) or immobilized (red) modes. At left, in the absence of netrin-1, the few immobilized DCC receptors form nanoclusters (yellow) lockdown within confined diffusion surface (light blue), the free diffusing DCC receptors hold dynamically exchange between free space and confined space. At right, in the presence of netrin-1, nanoclusters of immobilized DCC receptors grow (in size and area) by recruiting and stabilizing free diffusing receptors due to netrin-1 interactions mediated by netrin-1 glycosylation sites and, to some extent, netrin-1 interactions with other proteins (integrins). The growth of nanoclusters area within the confined diffusion surface produce decrease of confined diffusion surface.

Crystallographic studies have previously shown the existence of one interaction site in the LamVI domain and one in the EGF-3 domain (***Xu et al., 2014***), but that the three glycosylation sites of the LamVI domain did not point at the receptor (***Finci et al., 2014***). This could be due the lack of FN-4 in these experiments or to the limited glycosylation of the asparagine residues in crystalized complexes, due to their chemical and conformational heterogeneity (***Chang et al., 2007***). A partial glycan structure on these sites was identified as hetero-oligopolysaccharides of mannose and acetylglucosamine (***Grandin et al., 2016***), which correspond to only a small fraction of the full chain of glycosylation (***Geisbrecht et al., 2003***). The lack of fully glycosylated netrin-1 in a crystal complex with DCC has therefore prevented to implicate glycosylation in the interaction (***Krüger, 2014***). Here, our mutant approach shows that glycosylation at those sites is instrumental for netrin-1 capture by DCC-expressing cells, colocalization with DCC and the subsequent growth of immobilized DCC clusters (Figs 1-5). The abolition of netrin-1 effects by an antibody against DCC extracellular domain (Fig. 3) completes the picture by which the effects result from the direct interaction between glycosylated netrin-1 and DCC extracellular domain.

The C-terminus of netrin-1 promotes axon outgrowth in spinal cord explants (***Serafini et al., 1994***) and involves other interactions than with DCC (***Yebra et al., 2003; Stanco et al., 2009; Nikolopoulos and Gian 2005***). It is highly positively charged and has a strong affinity for cell membranes and the extracellular matrix (***Kappler et al., 2000; Tessier-Lavigne and Goodman, 1996***). Here, we demonstrate that netrin-1 lacking the C-terminal domain is captured by cells in proportion of their expression of DCC (Figs 2 and S1), supporting that this domain is not involved in the interaction with DCC. However, the lack of C-terminal domain in netrin-1 results in effects on DCC only different in magnitude compared to the lack of glycosylation (Figs 3-5). This supports that the immobilization and cluster growth of DCC that is promoted by glycosylation may in part be mediated by interactions involving the C-terminal domain and other proteins. The RGE mutant shows that impairment of interactions with integrins recapitulates most of the effects of the deletion of the whole C-terminal domain, to a milder extent in some cases (Figs 1-5). Therefore, interactions with integrins through an intact RGD site is likely to account for most of the interactions involving the C-terminal domain. This does not exclude other interactions such as with the extracellular matrix, and is consistent with an involvement of mechanical forces in netrin-1 signalization (***Stanco et al., 2009***).

Overall, our findings underscore that netrin-1 signaling through DCC operates through pre-existing platforms whose size and composition modifications are central elements of the signaling process. These modifications offer a promising array of regulation options, enriching our understanding of the intricate netrin-1 DCC signaling pathway.

## Methods and Materials

### Chimeric netrin and DCC constructs

The plasmid constructs encoding full-length netrin-1 (FL, Fig. 1A) and truncated netrin-1 lacking the C-domain, both fused to mCherry at the C terminus, were kindly provided by Simon W. Moore (Columbia University, New York) (***Moore et al., 2009***). These two constructs were modified in FL and dC by a fragment comprising a 6xHis tag and a SpeI recognizing site synthesized by GeneArt (Life Technologies) and inserted between the SgrAI and MfeI sites present at the C-terminal end of mCherry (restriction enzymes provided by NEB). Based on the FL plasmid, the constructs deleted of the LamVI domain (dLamVI) or deleted of the entire netrin sequence (mCherry) were amplified by PCR followed by homologous recombination according to the inFusion deletion protocol (Clontech; dLamVI primers: ATTGTAGAACGGTAGCCGCCCGTGCTCGTCGTAGCA and GCCGATGGCAAGAT-GTTACCCGTGCTCGTCGTAGCA ; mCherry primers: TGTCGAACGGTAGCCGCTGACGGTTCACTAAACC-AGCTCT and CGGCTACCGTTCGACATCGCCACCATGGTGAGC). GlycoM and RGE were generated by PCR of FL using oligonucleotides with mismatch according to inFusion point-mutation protocol (Clontech). For the glycoM plasmid, the four N-glycosylation sites were abolished by replacing asparagines (AAC) by alanines (GCC): from N-terminal to C-terminal : mutation ‘A’: GTGCCACCTCT-GCGCCGCCTCCGACCCC, mutation ‘B’: CTCAACAACCCGCACGCCCTGACGTGCTGGCA, mutation ‘C’: TGCAGTACCCGCACGCCGTCACCCTCACGC, mutation ‘D’: CCGGCCAAACCTGCGCCCAAACCACGGGGC (Fig. 2A). For the RGE plasmid, the RGD sequence (GAC) was replaced by the RGE sequence (GAG) using the GCTGCGGCGCGGGGAGCAGACCCTGTGGGTG primer. The DCC-GFP plasmid was a gift from Simon W. Moore (Columbia University, New York) (***Moore et al., 2009***). To perform super-resolution microscopy GFP was replaced by the photoactivatable GFP from the pPAGFP-N1 vector. The DCC-GFP D1290N mutant plasmid was generated by point mutation (GAC to AAC oligo CCAA-CACTCTCAGTGAACCGAGGTTTCGGAG ; Infusion, Clontech).

### Cell lines, culture and reagents

Caco-2 TC7 (gift from Françoise Poirier’s lab, Institut Jacques Monod) or HEK293T (ATCC Catalog No. CRL-11268) cell lines stably expressing mCherry-netrin-1 constructs or mCherry alone were generated by using TurboFect (Life Technologies) and maintained in a selective medium (DMEM Gibco + GlutaMAX + 4.5 g/l D-Glucose + Pyruvate (Life Technologies) + 10% Fetal Bovine Serum (Life Technologies) supplemented with 100 μg/ml G418 Geneticin (Life Technologies) during 2 weeks at 37 °C in a humidified 5% CO_2_ atmosphere. Fluorescent Caco-2 and HEK293T cells were sorted by Fluorescence Activated Cell Sorting (FACS) two times and kept under selection. Cells were imaged in non-fluorescent culture medium (FluoroBrite 4.5 g/l Glucose, 1% PenStrep, 2% FBS, L-Glutamine from Life Technologies). For secretion and capture, colocalization, FRAP, and SPT experiments, Caco-2 cells were transiently transfected with DCC-GFP or GFP plasmids, for PALM experiments HEK293T cells were transfected with DCC-PAGFP (TurboFect, Life Technologies). The HEK EBNA cell line stably expressing unlabeled recombinant netrin-1 (HEK NTN1) was generated previously (***Kennedy et al., 1994***). DCC monoclonal antibody (clone AF5, Thermo Fisher Scientific) from mouse was used for FRAP experiments.

### Chemotaxis assay for netrin-1-mCherry functionality

Aggregates of HEK293T stably expressing FL were co-cultured with rhombic lip explants from E12 mouse embryos embedded in rat-tail collagen according to the method described before (***Causeret et al., 2002***). The HEK NTN1 cell line stably expressing unlabeled recombinant netrin-1 and conventionally used for the study of axonal guidance, served as a positive control. After 72 hours in a 5% CO_2_, 37 °C, 95% humidity incubator, collagen assays were fixed in 4% paraformaldehyde. The axons were labeled with a monoclonal antibody against neuron-specific class III *β*-tubulin (clone Tuj-1 Babco) and the nuclei with DAPI (Sigma Aldrich). Pictures of explant were taken using a Leica fluorescent microscope and a Photometrics Coolsnap fx monochrome CCD camera (Roper Scientific, Duluth, GA). The blue channel allows visualization of cell nuclei staining with DAPI, the green channel allows visualization of Tuj1 staining with an Alexa-488-conjugated secondary antibody and the red channel allows visualization of expressed netrin-1-mCherry. Each explant was virtually divided into four quadrants with respect to the source of secreting netrin-1 from HEK cell aggregate. The axon outgrowth and nuclear migration effects are determined by measuring the area covered by Tuj1 immunostaining and by DAPI labeling, respectively, outside of the explant over proximal and distal quadrants from the netrin-1 expressing cell aggregates as indicated in Fig. 1A. The fluorescence associated with the body of the explant was excluded from the analysis.

### Netrin secretion assay by Fluorescence Activated Cell Sorting (FACS)

Caco-2 cells expressing DCC-GFP, GFP (“green cells”) or transfected with an empty vector (Ø) were co-cultured for 48h in normal culture conditions with Caco-2 cells expressing netrin-1-mCherry constructs (dC, dLamVI, RGE or glycoM), mCherry (“red cells”) or transfected with an empty vector (Ø). Cells were then treated with Accutase at 37 °C and filtered through 30 μm filters (CellTrics, Sysmex) to discard aggregates. Each condition contained about 90 000 cells and was repeated three times. Samples were analyzed with FACSAria Fusion flow cytometer (BD biosciences, Le Pont de Claix, France) equipped with two laser lines (488 and 561 nm). Data were acquired with BD CellQuest PRO software and analyzed with FlowJo software (Tree Star, Inc., Ashland, OR). To discriminate un-labelled cells (blue dots), DCC-GFP or GFP cells that did not capture netrin-1-mCherry (green dots), netrin-1-mCherry or mCherry cells (red dots) and DCC-GFP or GFP cells that captured mCherry(-netrin-1) (yellow dots), a sequential gating protocol was applied as follows:

1. Green threshold: First, on cells transfected with empty vectors, the cumulative intensity distribution in the red channel was fitted with a Gaussian distribution of mean *μ* and standard deviation *σ* to define a threshold *μ* + 3*σ* below which are 99.87% of those cells and above which cells were considered red. Next, on cells expressing mCherry(-netrin-1) constructs, the cumulative intensity distribution above this threshold was fitted with a Gaussian distribution to define a second threshold *μ* + 3*σ*, consistently close across all six mCherry(-netrin-1) constructs, above which cells were considered green.
2. Red threshold: On co-cultured cells, the cumulative intensity distributions above the green threshold (defined in step 1) were fitted with one or two Gaussian distributions. The best fit was scored with

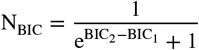

where BIC_1_ and BIC_2_ are the Bayesian information criterions for one and two Gaussians, respectively. N_BIC_ ranges from 0 (one Gaussian) to 1 (two Gaussians). When N_BIC_ > 0.5, the two populations were separated at the midpoint between their means, which defines the red threshold. Based on this threshold, the proportion *ρ*_*D*_ of DCC-GFP or GFP cells that captured mCherry(-netrin-1) (yellow dots) over all DCC-GFP or GFP cells (green and yellow dots) was determined for every co-culture.

Finally, we assessed the correlation between the expression levels of DCC-GFP or GFP and the capture of mCherry(-netrin-1) with the Pearson’s correlation coefficient r of green and red intensities on all DCC-GFP or GFP cells (green and yellow dots).

### Colocalization analyses by confocal imaging

Cells were imaged in glass bottom dishes (35 mm dish, 10 mm glass diameter, MatTek) with a Zeiss LSM780 confocal microscope equipped with a Plan-Apochromat 63x/1.4 oil objective. Images of DCC-GFP (488-nm excitation) and of mCherry(-netrin-1) σ 61-nm excitation) were recorded sequentially with Zen software (Zeiss). Correlation analysis was performed using FIJI. The plasma membrane region of interest (ROI_memb_, Fig. 2C in cyan) was obtained as follows: (1) images were blurred to attenuate the noise, (2) cell were then segmented by applying a manual threshold (outer boundaries of ROI_memb_) and (3) these cell boundaries were eroded of 10 pixels and manually adjusted to include all protrusions (inner boundaries of ROI_memb_). The correlation between red and green intensities within ROI_memb_ was assessed with the Pearson’s correlation coefficient r.

### FRAP experiments and analysis

Cells were imaged on a Leica DMI6000 inverted microscope equipped with a Yogogawa CSU22 spinning-disk head, a Plan-Apochromat 100x/1.4 oil-immersion objective and an EMCCD camera (Photometrics QuantEM 512S). Six spots of 2×2 pixels on the basal membrane were photo-bleached at 100% laser power of a 473-nm diode laser, and image acquisitions were performed with a 491-nm diode laser every 1 s for 5 s prebleach, then every 0.5 s for 30 s and 5 s for 300 s after bleach. For each cell, fluorescence levels were analyzed with ImageJ in 8 ROIs of the same surface (“Base”: background, “Whole”: not bleached area, 6 bleached spots). For each ROI, the intensity I was measured and normalized such that *I*_Base_(*t*) = 0% and *I*_ROI_(pre − bleach) = 100%. Normalized FRAP timeseries were pooled and mean curves were fitted by adapting the reduced two-state model from (***Sprague et al., 2004***) to account for the processes of binding and diffusion of DCC, leading to the following reaction-diffusion model :

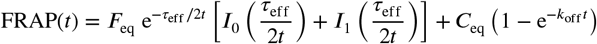

where *F*_eq_ and *C*_eq_ are the diffusive and reactive fractions, *I*_0_ and *I*_1_ are modified Bessel functions of the first kind, *k*_off_ is the off-rate of the reactive fraction, and *τ*_eff_ is defined as:

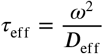

with *w*^2^ the bleached area and *D*_eff_ the effective diffusion coefficient. The immobile fraction is defined by the following expression:

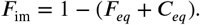

### Single Particle Tracking (SPT) and analysis

Imaging was performed on a TIRF motorized microscope (ELYRA Zeiss) equipped with a Plan-Apochromat 100x/1.46 oil objective and an EMCCD camera (Andor, iXon Ultra 897; frame-transfer mode; vertical shift speed: 0.9 μs; -70°C), using a 488 nm excitation laser and an exposure time of 50 ms. The incubation chamber maintained a humidified 37 °C atmosphere with 5% CO_2_. SPT data was analyzed (detection and tracking) and converted into trajectories using a custom-written Matlab (MathWorks Inc.) implementation of the Multiple-Target Tracing (MTT) algorithm (***Sergé et al., 2008***). To assess whether tracked particles are single, we developed a custom MATLAB (Mathworks, Natick, MA) script to measure the temporal decay of particles fluorescence intensity (from MTT). Particles were considered single when their decay displayed a single step to reach average local background intensity as defined within a ROI of 20×20 pixels centred on the particle position or on its last known position after it disappears. The percentage of single particles in 1000 randomly picked trajectories was around 50% for all conditions. The SPT analysis was then carried out exclusively on trajectories that were identified as corresponding to single entities. Any trajectories that did not meet this criterion were excluded and discarded from further analysis. First the mean square displacement (MSD) was computed up to 12 time-lags for trajectories comprising at least 20 positions according to:

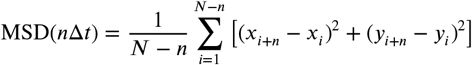

where *x*_*i*_ and *y*_*i*_ are the coordinates of a particle in the *i*^th^ image, *N* is the total number of images of the experiment and Δ*t* is the time lag between two images. The instantaneous diffusion coefficient *D*_inst_ was measured by fitting each trajectory with MSD = *D*_*inst*_*t* over the first three intervals of each trajectory. Using the average resolution of each particle provided by MTT, we determine a threshold corresponding to the smallest measurable diffusion coefficient:

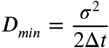

where *σ* the mean localization error in spot detection. Particles exhibiting *D*_*inst*_ < *D*_*min*_ were then considered as immobile. The remaining MSD curves were then fitted with a model of confined diffusion:

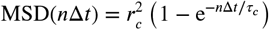

where 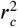 is the confinement area and τ _*c*_ the characteristic time to reach 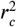. We used *τ*_*c*_ to sort particle in sub-diffusion (τ _*c*_ < 200 ms) and free diffusion categories. Finally, MSD curves were averaged according to their diffusion type: ‘Immobile’, ‘Confined’ and ‘Free’.

### Validation of SPT analyses

In Matlab 9.4 (Mathworks, Natick, MA), we simulated a SPT dataset of 18 000 trajectories using a Monte Carlo algorithm with a computational time step of 20 Hz within 256 × 256 pixels of 100 nm^2^ and three diffusion modes (6000 trajectories each) (Fig. S2):

1. Simple diffusion mode: Brownian diffusion simulations were performed by generating a set of random displacements at each time step as explained in ***Michalet*** (***2010***). These displacements follow a zero-mean Gaussian distribution with a standard deviation equal to (2*D*Δ*t*)^1/2^, where *D* is the diffusion coefficient and Δ*t* is the time step between two successive frames.
2. Confined diffusion mode: particles were initially placed at random positions defined as the center of a the 2D circular boundary of radius *r*_*c*_. At each time step, *x* and *y* displacements from simple diffusion were sampled from a Gaussian distribution with zero mean and standard deviation (2*D*Δ*t*)^1/2^ (mode (1)). The position of the particles was then updated only if the distance from the starting point to the new coordinates is smaller than *r*_*c*_.
3. Immobile mode: particles follow a simple diffusion mode but with a diffusion coefficient *D*_*inst*_ randomly chosen for each trajectory between [*D*/10, *D*/5], where *D* is diffusion coefficient used in mode (1).

To simulate localization noise, we added to each position a normal Gaussian noise with zero mean and a standard deviation set arbitrarily to 10 × *D*Δ*t*. The diffusion coefficient *D* = 0.04 × 10^−12^ μm^2^/s and radius of confinement *r*_*c*_ = 225 nm were taken from the experimental data. Results of the classification of simulations are summarized in the confusion matrix (Fig. S2.C). The precision (true positives/all positives) and recall (true positive/(true positive/false positive)) of this classification were above 80% for all modes. Its overall accuracy (true positives and negatives/all trajectories) was above 85%.

### PALM experiments and nanoclusters analysis

PALM imaging was performed on cells fixed in paraformaldehyde on an ELYRA Zeiss microscope equipped with a Plan-Apochromat 100x/1.46 oil objective in total internal reflection fluorescence (TIRF) mode, using the same laser wavelength 488 nm for simultaneous activation and excitation of PAGFP (laser intensity set to 10 μW measured at the rear aperture of the objective). Fluorescence emission was detected with an EMCCD camera (Andor, iXon Ultra 897) and acquired with Zen black 2012 (Zeiss). Time lapses of 2000 frames of 25 μm × 25 μm were collected with an exposure time of 50 ms per frame and with drift correction. Nanocluster analyses were performed on cropped images of 25 μm^2^. We used the SLIMfast MATLAB (MathWorks) code (Institut Curie, France) (***Normanno et al., 2015***) to recover single molecule localizations. The average localization precision ranged from 18 to 32 nm (half width at half height of gaussian fit). To remove blinking (and sequential localization of the same molecule), we used the Distance Distribution Correction (DDC) algorithm (***Bohrer et al., 2021***). Briefly, the duration over which a fluorophore blinks is first obtained from the rate at which the pairwise distance distribution converges to a steady state. Here, the Z parameter (normalized, summed differences of the cumulative distributions for each frame interval) reached a plateau for *N* = 200 frames. This value of *N* and distance *ε* = 60 nm (two times the precision of localization) were then fed in the DDC algorithm to remove multiple detections of blinking fluorophores. We then used DBSCAN (***Tran et al., 2013***) to identify and quantify nanoclusters with a minimal cardinality *k* = 4 within a distance *ε* = 60 nm among remaining localizations. We finally retrieved the number of clusters, their cardinality and surface area, and the fraction of localization in clusters all as a function of localization number for each field of view.

### Hierarchical Cluster Analysis (HCA)

Hierarchicalclustering of netrin-1 constructs was performed with the Python library pvclust (***Suzuki and Shim 2006***) using the Euclidean distance metric and the Ward clustering method on a dataset of 9 metrics obtained from FRAP (immobile fraction, dissociation rate), SPT (immobile fraction, diffusion coefficient, confinement size), colocalization (Pearson coefficient), or nanocluster analyses (Cluster area, % in clusters and number of molecules in clusters as a function of localization number). To balance statistical weight between metrics and maximize bootstrap resampling power, a linear transformation normalized each metric value between the mCherry condition set to 0 and the FL condition set to 1. The statistical significance of each cluster was measured by two types of p-values calculated by bootstrap resampling: the Bootstrap Probability value (BP) and the Approximately Unbiased (AU) p-value.

### Other statistical analyses

Data were analyzed with GraphPad Prism software (GraphPad Software, Inc., San Diego, CA, USA). All values are expressed as mean ± SE. Statistical significance was assessed with Student’s paired t-test or two-way ANOVA. Resulting *P* values are ns when *P* > 0.05; * when *P* ≤ 0.05, ** when *P* ≤ 0.01, *** when *P* ≤ 0.001 and **** when *P* < 0.0001.

## Supporting information

Latex file with cls, bib and images

## Acknowledgments

We especially thank Simon W. Moore (Columbia University, New York, USA) and Françoise Poirier’s lab (Institut Jacques Monod, Paris, France) for sharing reagents, Mathieu Coppey (Institut Curie, Paris, France) and Nicolas Moisan (Institut Jacques Monod, Paris, France) for their technical help with the PALM analysis. This work was supported by the Centre National de la Recherche Scientifique (CNRS) and France-BioImaging (ANR-10-INSB-04). KU had a fellowship from the company Carl Zeiss SAS (CIFRE nº 2012/1059). PPG have received funding from the German Academic Exchange Service (DAAD). We acknowledge the ImagoSeine core facility of the Institut Jacques Monod, member of the Infrastructures en Biologie Santé et Agronomie (IBiSA) and France-BioImaging (ANR-10-INSB-04) infrastructures.

**Figure S1.**
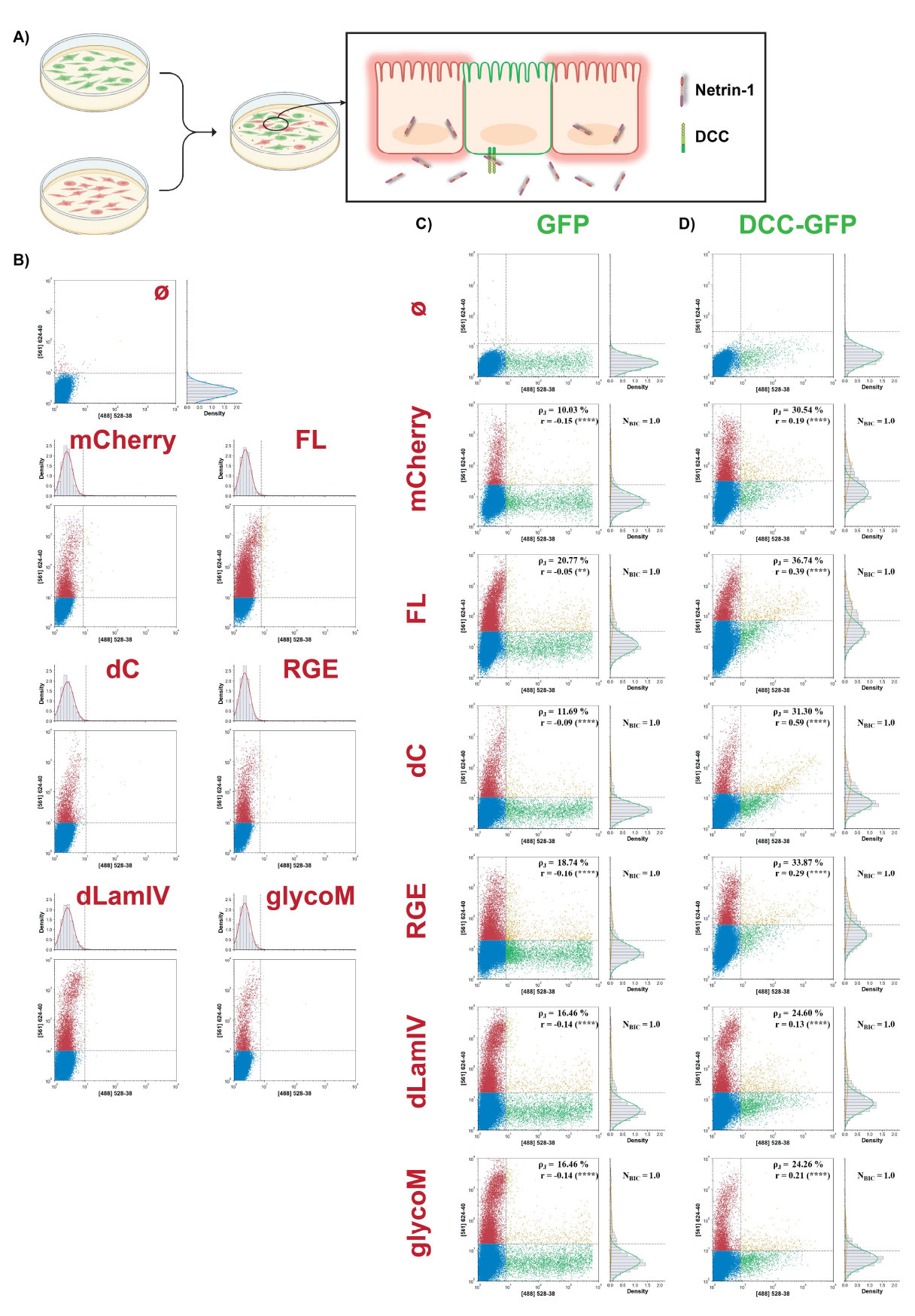
(A) Principle of the co-culture between Caco-2 cells expressing mCherry or netrin-1 constructs (red) and Caco-2 cells expressing GFP or DCC-GFP (green). (B) Cytograms of co-cultures between Caco-2 cells transfected with an empty vector (Ø), or expressing mCherry, FL, dC, RGE, dLamVI or glycoM, and Caco-2 cells transfected with an empty vector (Ø), which shows cells expressing no construct (blue dots) and cells expressing mCherry or netrin-1 constructs (red dots). (C) Cytograms of co-cultures between Caco-2 cells transfected with an empty vector (Ø), or expressing mCherry, FL, dC, RGE, dLamVI or glycoM, and Caco-2 cells transfected with GFP or DCC-GFP, which shows cells expressing no construct (blue dots), cells expressing mCherry or netrin-1 constructs (red dots), the original DCC-GFP cells (green dots), and DCC-GFP cells that captured netrin-1 constructs (yellow dots).

**Figure S2.**
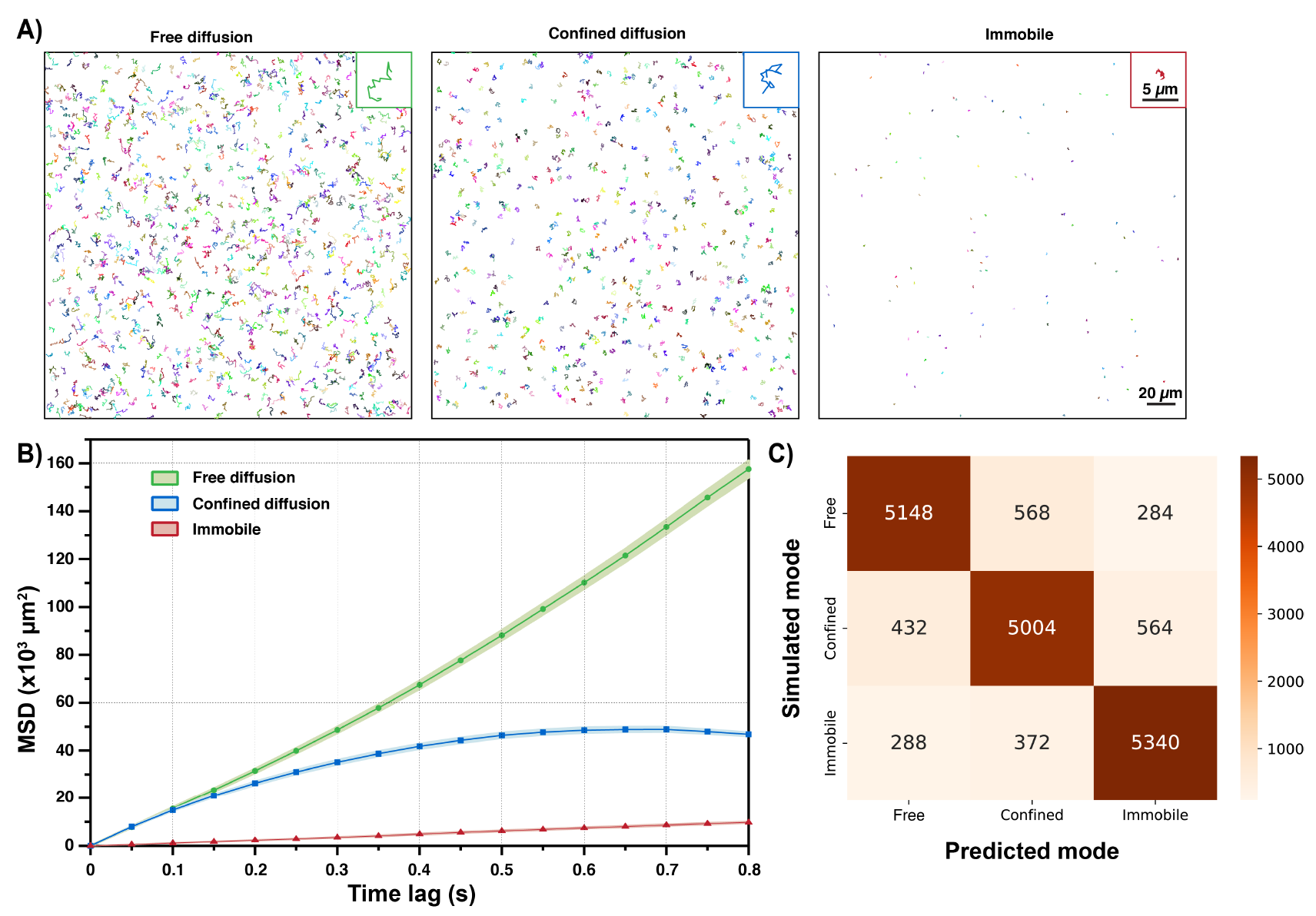
SPT. (A) Typical individual trajectories simulated for each diffusion mode: free diffusion, confined diffusion and immobile. (B) MSD curves for the trajectories shown in (A). (C) Confusion matrix of the diffusion mode classification (*n* = 18 000 trajectories).

**Figure S3.**
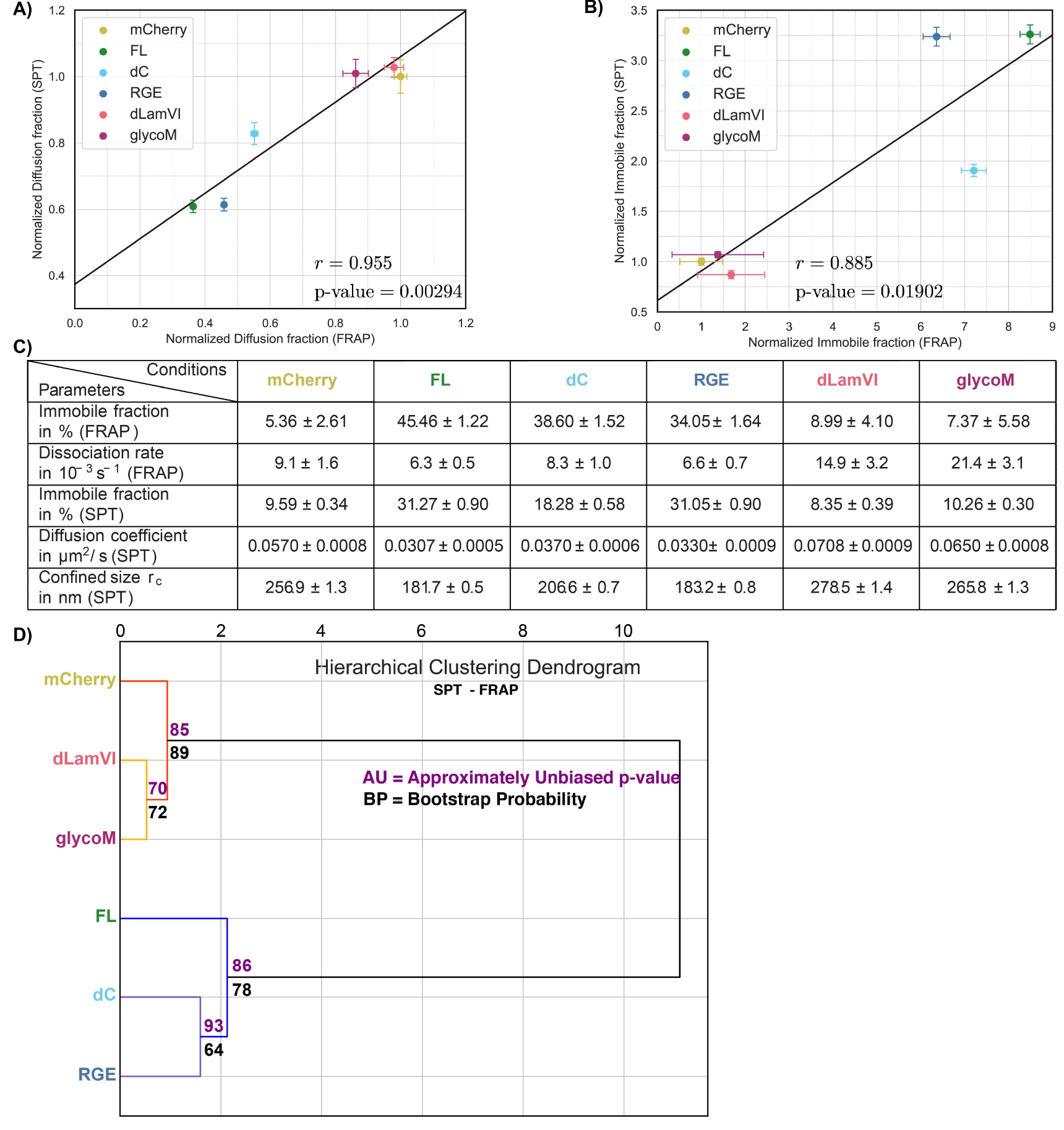
Correlation SPT-FRAP. Correlation of normalized diffusion fractions (A) and immobile fractions (B) between SPT and FRAP experiments for netrin-1 variants (FL, dC, RGE, dLamVI and glycoM) and mCherry. r: Pearson’s correlation coefficient. (C) FRAP and SPT metrics used for hierarchical clustering in (D). (D) Hierarchical clustering of mCherry and netrin-1 constructs based on FRAP and SPT data.

**Figure S4.**
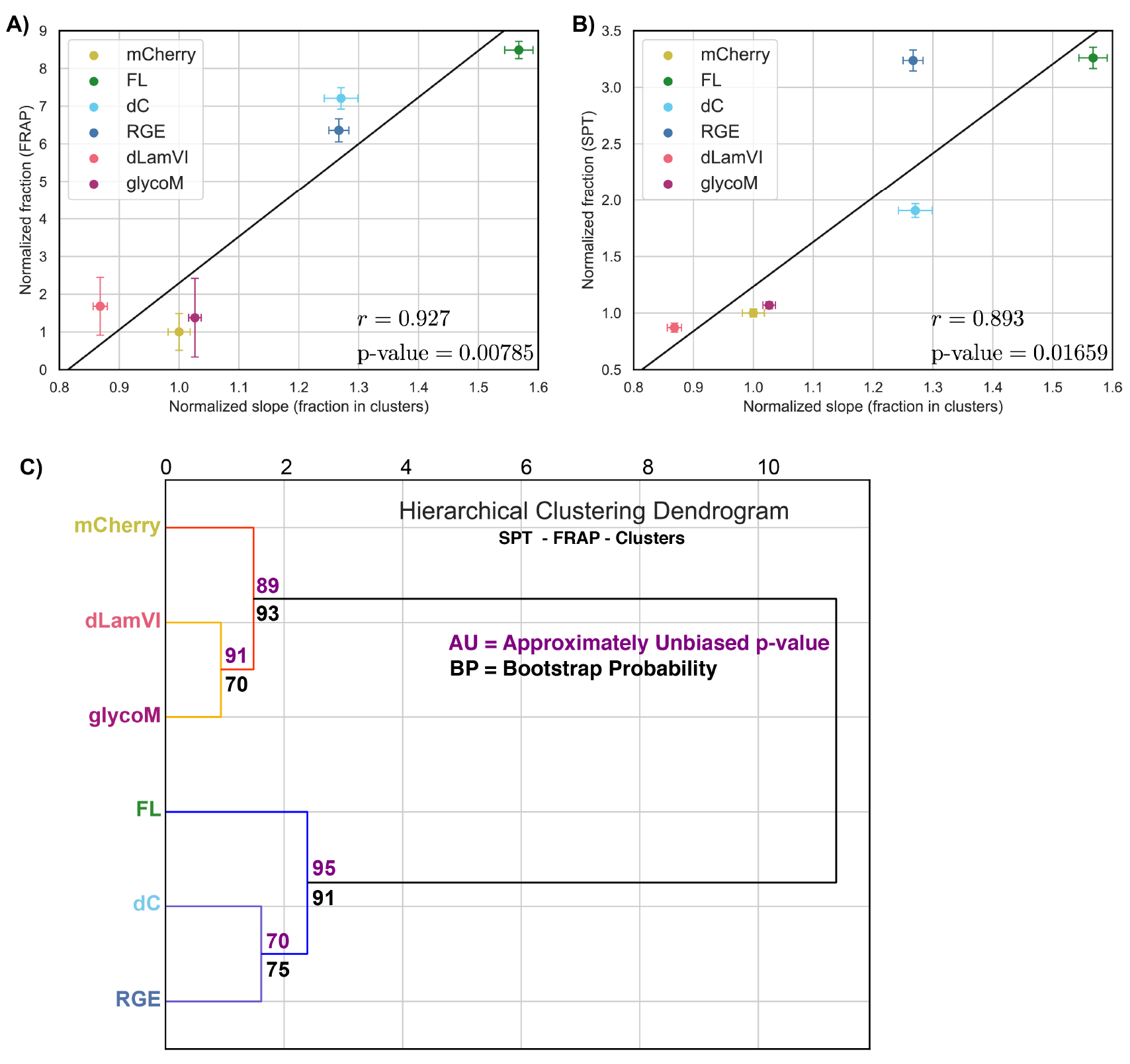
Correlation FRAP-SPT-CIusters. Normalized immobile fractions from FRAP (A) and SPT (B) vs. normalized fraction in cluster per localization (see Fig. 5E) for mCherry, FL, dC, RGE, dLamVI and glycoM, r: Pearson’s correlation coefficient. (C) Hierarchical clustering of mCherry and netrin-1 constructs based on FRAP (immobile fraction, dissociation rate), SPT (immobile fraction, diffusion coefficient, confinement size), colocalization (Pearson coefficient), and nanocluster analyses (Cluster area, % in clusters and number of molecules in clusters as a function of localization number).

## Notes

### Competing Interest Statement

The authors have declared no competing interest.

### Summary of Updates

This version of the manuscript has been compiled with the elife Latex class and the typographical errors have been corrected

## References

Baker, K. A., Moore, S. W., Jarjour, A. A., and Kennedy, T. E. (2006). When a diffusible axon guidance cue stops diffusing: roles for netrins in adhesion and morphogenesis. Curr. Opin. Neurobiol., 16(5):529–34.

Bányai, L. and Patthy, L. (1999). The NTR module: Domains of netrins, secreted frizzled related proteins, and type I procollagen C-proteinase enhancer protein are homologous with tissue inhibitors of metalloproteases. Protein Sci., 8(8):1636–1642.

Bloch-Gallego, E., Ezan, F., Tessier-Lavigne, M., and Sotelo, C. (1999). Floor plate and netrin-1 are involved in the migration and survival of inferior olivary neurons. J. Neurosci., 19(11):4407–4420.

Bohrer, C. H., Yang, X., Thakur, S., Weng, X., Tenner, B., McQuillen, R., Ross, B., Wooten, M., Chen, X., Zhang, J., Roberts, E., Lakadamyali, M., and Xiao, J. (2021). A pairwise distance distribution correction (DDC) algorithm to eliminate blinking-caused artifacts in SMLM. Nat. Methods, 18(6):669–677.

Bouchard, J. F., Moore, S. W., Tritsch, N. X., Roux, P. P., Shekarabi, M., Barker, P. A., and Kennedy, T. E. (2004). Protein Kinase A Activation Promotes Plasma Membrane Insertion of DCC from An Intracellular Pool: A Novel Mechanism Regulating Commissural Axon Extension. J. Neurosci., 24(12):3040–3050.

Causeret, F., Danne, F., Ezan, F., Sotelo, C., and Bloch-Gallego, E. (2002). Slit antagonizes netrin-1 attractive effects during the migration of inferior olivary neurons. Dev. Biol., 246(2):429–440.

Chang, V. T., Crispin, M., Aricescu, A. R., Harvey, D. J., Nettleship, J. E., Fennelly, J. A., Yu, C., Boles, K. S., Evans, E. J., Stuart, D. I., Dwek, R. A., Jones, E. Y., Owens, R. J., and Davis, S. J. (2007). Glycoprotein structural genomics: solving the glycosylation problem. Structure (London, England : 1993), 15(3):267–273.

Chen, J. Y., He, X. X., Ma, C., Wu, X. M., Wan, X. L., Xing, Z. K., Pei, Q. Q., Dong, X. P., Liu, D. X., Xiong, W. C., and Zhu, X. J. (2017). Netrin-1 promotes glioma growth by activating NF-,cB via UNC5A. Sci. Rep., 7(1).

Dent, E. W., Barnes, A. M., Tang, F., and Kalil, K. (2004). Netrin-1 and semaphorin 3a promote or inhibit cortical axon branching, respectively, by reorganization of the cytoskeleton. J. Neurosci., 24(12):3002–3012.

Dun, X. P. and Parkinson, D. B. (2017). Role of netrin-1 signaling in nerve regeneration. Int. J. Mol. Sci., 18(2):491.

Fazeli, A., Dickinson, S. L., Hermiston, M. L., Tighe, R. V., Steen, R. G., Small, C. G., E T Stoeckli, K. K.-M., Masu, M., Rayburn, H., Simons, J., Bronson, R. T., Gordon, J. I., Tessier-Lavigne, M., and Weinberg, R. A. (1997). Phenotype of mice lacking functional deleted in colorectal cancer (dcc) gene. Nature, 386(6627):796–804.

Fearon, E. R. and Vogelstein, B. (1990). A genetic model for colorectal tumorigenesis. Cell, 61(5):759–767.

Finci, L., Zhang, Y., Meijers, R., and Wang, J.-H. (2015). Signaling mechanism of the netrin-1 receptor DCC in axon guidance. Prog. Biophys. Mol. Biol., 118(3):153–160.

Finci, L. I., Krüger, N., Sun, X., Zhang, J., Chegkazi, M., Wu, Y., Schenk, G., Mertens, H. D., Svergun, D. I., Zhang, Y., huai Wang, J., and Meijers, R. (2014). The Crystal Structure of Netrin-1 in Complex with DCC Reveals the Bifunctionality of Netrin-1 As a Guidance Cue. Neuron, 83(4):839–849.

Fitamant, J., Guenebeaud, C., Coissieux, M. M., Guix, C., Treilleux, I., Scoazec, J. Y., Bachelot, T., Bernet, A., and Mehlen, P. (2008). Netrin-1 expression confers a selective advantage for tumor cell survival in metastatic breast cancer. Proc. Natl. Acad. Sci. U.S.A., 105(12):4850–4855.

Forcet, C., Ye, X., Granger, L., Corset, V., Shin, H., Bredesen, D. E., and Mehlen, P. (2001). The dependence receptor DCC (deleted in colorectal cancer) de?nes an alternative mechanism for caspase activation. Proc. Natl. Acad. Sci. U.S.A.

Geisbrecht, B. V., Dowd, K. A., Barfield, R. W., Longo, P. A., and Leahy, D. J. (2003). Netrin binds discrete subdomains of DCC and UNC5 and mediates interactions between DCC and heparin. J. Biol. Chem., 278(35):32561– 32568.

Gopal, A. A., Rappaz, B., Rouger, V., Martyn, I. B., Dahlberg, P. D., Meland, R. J., Beamish, I. V., Kennedy, T. E., and Wiseman, P. W. (2016). Netrin-1-Regulated Distribution of UNC5B and DCC in Live Cells Revealed by TICCS. Biophys. J., 110(3):623–634.

Grandin, M., Meier, M., Delcros, J. G., Nikodemus, D., Reuten, R., Patel, T. R., Goldschneider, D., Orriss, G., Krahn, N., Boussouar, A., Abes, R., Dean, Y., Neves, D., Bernet, A., Depil, S., Schneiders, F., Poole, K., Dante, R., Koch, M., Mehlen, P., and Stetefeld, J. (2016). Structural Decoding of the Netrin-1/UNC5 Interaction and its Therapeutical Implications in Cancers. Cancer Cell, 29(2):173–185.

Guirland, C., Suzuki, S., Kojima, M., Lu, B., and Zheng, J. Q. (2004). Lipid rafts mediate chemotropic guidance of nerve growth cones. Neuron, 42(1):51–62.

Hamelin, M., Zhou, Y., Su, M. W., Scott, I. M., and Culotti, J. G. (1993). Expression of the unc-5 guidance receptor in the touch neurons of c. elegans steers their axons dorsally. Nature, 364(6435):327–30.

Hebrok, M. and Reichardt, L. F. (2004). Brain meets pancreas: netrin, an axon guidance molecule, controls epithelial cell migration. Trends Cell Biol., 14(4):153–5.

Hedgecock, E. M., Culotti, J. G., and Hall, D. H. (1990). The unc-5, unc-6, and unc-40 genes guide circumferential migrations of pioneer axons and mesodermal cells on the epidermis in c. elegans. Neuron., 4(11):61–85.

Hedrick, L., Cho, K. R., Fearon, E. R., Wu, T. C., Kinzler, K. W., and Vogelstein, B. (1994). The DCC gene product in cellular differentiation and colorectal tumorigenesis. Genes Dev., 8(10):1174–1183.

Hérincs, Z., Corset, V., Cahuzac, N., Furne, C., Castellani, V., Hueber, A.-O., and Mehlen, P. (2005). DCC association with lipid rafts is required for netrin-1-mediated axon guidance. J. Cell Sci., 118(Pt 8):1687–92.

Hinck, L. (2004). The versatile roles of “axon guidance” cues in tissue morphogenesis. Dev. Cell, 7(6):783–93.

Kappler, J., Franken, S., Junghans, U., Hoffmann, R., Linke, T., Müller, H. W., and Koch, K. W. (2000). Glycosaminoglycan-binding properties and secondary structure of the C-terminus of netrin-1. Biochem. Biophys. Res. Commun., 271(2):287–291.

Keino-Masu, K., M Masu, L. H., Leonardo, E. D., Chan, S. S., Culotti, J. G., and Tessier-Lavigne, M. (1996). Deleted in colorectal cancer (dcc) encodes a netrin receptor. Cell, 87(2):175–185.

Kennedy, T. E., Serafini, T., de la Torre, J. R., and Tessier-Lavigne, M. (1994). Netrins are diffusible chemotropic factors for commissural axons in the embryonic spinal cord. Cell.

Kolodziej, P. A., Timpe, L. C., Mitchell, K. J., Fried, S. R., Goodman, C. S., Jan, L. Y., and Jan, Y. N. (1996). frazzled encodes a drosophila member of the dcc immunoglobulin subfamily and is required for cns and motor axon guidance. Cell, 87(2):197–204.

Krüger, N. (2014). Structural and functional analysis of the guidance cue molecule netrin-1 in complex with its receptor DCC unravels the molecular mechanisms of interaction. PhD thesis, Staats-und Universitätsbibliothek Hamburg Carl von Ossietzky.

Lim, Y.-s. and Wadsworth, W. G. (2002). Identi?cation of domains of netrin UNC-6 that mediate attractive and repulsive guidance and responses from cells and growth cones. J. Neurosci., 22(16):7080–7087.

Marcos, S., Backer, S., Causeret, F., Tessier-Lavigne, M., and Bloch-Gallego, E. (2009). Differential roles of Netrin-1 and its receptor DCC in inferior olivary neuron migration. Mol. Cell. Neurosci., 41(4):429–439.

Matsumoto, H. and Nagashima, M. (2010). Netrin-1 elevates the level and induces cluster formation of its receptor DCC at the surface of cortical axon shafts in an exocytosis-dependent manner. Neurosci. Res., 67(2):99– 107.

Mehlen, P., Rabizadeh, S., Snipas, S. J., Assa-Munt, N., Salvesen, G. S., and Bredesen, D. E. (1998). The DCC gene product induces apoptosis by a mechanism requiring receptor proteolysis. Nature, 395(6704):801–804.

Michalet, X. (2010). Mean square displacement analysis of single-particle trajectories with localization error: Brownian motion in an isotropic medium. Phys. Rev. E., 82(4 Pt 1):41914.

Moore, S. W., Biais, N., and Sheetz, M. P. (2009). Traction on immobilized netrin-1 is su?cient to reorient axons. Science, 325(5937):166.

Nikolopoulos, S. N. and Giancotti, F. G. (2005). Netrin-integrin signaling in epithelial morphogenesis, axon guidance and vascular patterning. Cell Cycle, 4(3):e131–5.

Normanno, D., Boudarène, L., Dugast-Darzacq, C., Chen, J., Richter, C., Proux, F., Bénichou, O., Voituriez, R., Darzacq, X., and Dahan, M. (2015). Probing the target search of DNA-binding proteins in mammalian cells using TetR as model searcher. Nat. Commun., 6:7357.

Petrie, R. J., Zhao, B., Bedford, F., and Lamarche-Vane, N. (2009). Compartmentalized DCC signalling is distinct from DCC localized to lipid rafts. Biol. Cell, 101(2):77–90.

Rappaz, B., Lai Wing Sun, K., Correia, J. P., Wiseman, P. W., and Kennedy, T. E. (2016). FLIM FRET Visualization of Cdc42 Activation by Netrin-1 in Embryonic Spinal Commissural Neuron Growth Cones. PloS one, 11(8):e0159405.

Serafini, T., Colamarino, S. A., Leonardo, E. D., H Wang, R. B., Skarnes, W. C., and Tessier-Lavigne, M. (1996). Netrin-1 is required for commissural axon guidance in the developing vertebrate nervous system. Cell, 87(6):1001–1014.

Serafini, T., Kennedy, T. E., Galko, M. J., Mirzayan, C., Jessell, T. M., and Tessier-Lavigne, M. (1994). The netrins de?ne a family of axon outgrowth-promoting proteins homologous to C. elegans UNC-6. Cell, 78(3):409–424.

Sergé, A., Bertaux, N., Rigneault, H., and Marguet, D. (2008). Dynamic multiple-target tracing to probe spatiotemporal cartography of cell membranes. Nat. Methods.

Sprague, B. L., Pego, R. L., Stavreva, D. A., and McNally, J. G. (2004). Analysis of binding reactions by ?uorescence recovery after photobleaching. Biophys. J., 86(6):3473–3495.

Stanco, A., Szekeres, C., Patel, N., Rao, S., Campbell, K., Kreidberg, J. A., Polleux, F., and Anton, E. S. (2009). Netrin-1-alpha3beta1 integrin interactions regulate the migration of interneurons through the cortical marginal zone. Proc. Natl. Acad. Sci. U.S.A., 106(18):7595–600.

Suzuki, R. and Shimodaira, H. (2006). Pvclust: an R package for assessing the uncertainty in hierarchical clustering. Bioinformatics (Oxford, England), 22(12):1540–1542.

Tessier-Lavigne, M. and Goodman, C. S. (1996). The molecular biology of axon guidance. Science (New York, N.Y.), 274(5290):1123–1133.

Tessier-Lavigne, M., Placzek, M., Lumsden, A. G. S., Dodd, J., and Jessell, T. M. (1988). Chemotropic guidance of developing axons in the mammalian central nervous system. Nature, 336(6201):775–8.

Tran, T. N., Drab, K., and Daszykowski, M. (2013). Revised DBSCAN algorithm to cluster data with dense adjacent clusters. Chemom. Intell. Lab. Syst., 120:92–96.

Xu, B., Goldman, J. S., Rymar, V. V., Forget, C., Lo, P. S., Bull, S. J., Vereker, E., Barker, P. A., Trudeau, L. E., Sadikot, A. F., and Kennedy, T. E. (2004). Critical roles for the netrin receptor deleted in colorectal cancer in dopaminergic neuronal precursor migration, axon guidance, and axon arborization. Neuroscience, 169(2):932–49.

Xu, K., Wu, Z., Renier, N., Antipenko, A., Tzvetkova-Robev, D., Xu, Y., Minchenko, M., Nardi-Dei, V., Rajashankar, K. R., Himanen, J., Tessier-Lavigne, M., and Nikolov, D. B. (2014). Structures of netrin-1 bound to two receptors provide insight into its axon guidance mechanism. Science, 344(6189):1275–1279.

Yamagishi, S., Yamada, K., Sawada, M., Nakano, S., Mori, N., Sawamoto, K., and Sato, K. (2015). Netrin-5 is highly expressed in neurogenic regions of the adult brain. Front. Cell. Neurosci., 9(APR).

Yebra, M., Montgomery, A. M., Diaferia, G. R., Kaido, T., Silletti, S., Perez, B., Just, M. L., Hildbrand, S., Hurford, R., Florkiewicz, E., Tessier-Lavigne, M., and Cirulli, V. (2003). Recognition of the neural chemoattractant netrin-1 by integrins a6p4 and a3p1 regulates epithelial cell adhesion and migration. Dev. Cell, 5(5):695–707.

Ylivinkka, I., Keski-Oja, J., and Hyytiäinen, M. (2016). Netrin-1: A regulator of cancer cell motility? European Journal of Cell Biology, 95(11):513–520. Integrated mechano-chemical signals in invasion.

