## Supplementary material for "Interactions of netrin-1 through its glycosylation sites immobilize Deleted in Colorectal Cancer (DCC) by favoring its constitutive clustering": Latex file with cls, bib and images: KarenV2.bib

@ARTICLE{Bloch1999,
  author = {E. Bloch-Gallego and F. Ezan and M. Tessier-Lavigne and  C. Sotelo},
  title = {Floor plate and netrin-1 are involved in the migration and survival of inferior olivary neurons},
  journal = {J. Neurosci.},
  year = {1999},
  volume = {19},
  pages = {4407--4420},
  number = {11},
  doi = {10.1523/JNEUROSCI.19-11-04407.1999},
}

@ARTICLE{Hedgecock1990,
  author = {E. M. Hedgecock and  J. G. Culotti and  D. H. Hall},
  title = {The unc-5, unc-6, and unc-40 genes guide circumferential migrations of pioneer axons and mesodermal cells on the epidermis in C. elegans},
  journal = {Neuron.},
  year = {1990},
  volume = {4},
  pages = {61--85},
  number = {11},
  doi = {10.1016/0896-6273(90)90444-k},
}


@ARTICLE{Serafini1996,
  author = {T Serafini and S A Colamarino and  E D Leonardo and  H Wang, R Beddington and W C Skarnes and  M Tessier-Lavigne},
  title = {Netrin-1 is required for commissural axon guidance in the developing vertebrate nervous system},
  journal = {Cell},
  year = {1996},
  volume = {87},
  pages = {1001--1014},
  number = {6},
  doi = {10.1016/s0092-8674(00)81795-x},
}

@ARTICLE{Keino1996,
  author = {K Keino-Masu  and  M Masu, L Hinck and  E D Leonardo and S S Chan and  J G Culotti and  M Tessier-Lavigne},
  title = {Deleted in Colorectal Cancer (DCC) encodes a netrin receptor.},
  journal = {Cell},
  year = {1996},
  volume = {87},
  pages = {175--185},
  number = {2},
  doi = {10.1016/s0092-8674(00)81336-7},
}

@ARTICLE{Kolodziej1996,
  author = {P A Kolodziej  and L C Timpe and  K J Mitchell and  S R Fried and  C S Goodman and L Y Jan and Y N Jan},
  title = {frazzled encodes a Drosophila member of the DCC immunoglobulin subfamily and is required for CNS and motor axon guidance},
  journal = {Cell},
  year = {1996},
  volume = {87},
  pages = {197--204},
  number = {2},
  doi = {10.1016/s0092-8674(00)81338-0},
}


@ARTICLE{Fazeli1997,
  author = {A Fazeli and S L Dickinson and M L Hermiston and R V Tighe and R G Steen and C G Small and E T Stoeckli, K Keino-Masu and M Masu and H Rayburn and J Simons and R T Bronson and J I Gordon and M Tessier-Lavigne and R A Weinberg},
  title = {Phenotype of mice lacking functional Deleted in colorectal cancer (Dcc) gene},
  journal = {Nature},
  year = {1997},
  volume = {386},
  pages = {796--804},
  number = {6627},
  doi = {10.1038/386796a0},
}

@ARTICLE{Tessier1988,
  author = {M. Tessier-Lavigne and M. Placzek and  A. G. S. Lumsden and  J. Dodd and  T. M. Jessell},
  title = {Chemotropic guidance of developing axons in the mammalian central nervous system},
  journal = {Nature},
  year = {1988},
  volume = {336},
  pages = {775--8},
  number = {6201},
  doi = {10.1038/336775a0},
}

@ARTICLE{Hamelin1993,
  author = {M. Hamelin and Y. Zhou and M. W. Su and  I. M. Scott and J. G. Culotti},
  title = {Expression of the UNC-5 guidance receptor in the touch neurons of C. elegans steers their axons dorsally},
  journal = {Nature},
  year = {1993},
  volume = {364},
  pages = {327--30},
  number = {6435},
  doi = {10.1038/364327a0},
}

@ARTICLE{Dent2004,
  author = {E. W. Dent and A. M. Barnes and F. Tang and K. Kalil},
  title = {Netrin-1 and Semaphorin 3A Promote or Inhibit Cortical Axon Branching, Respectively, by Reorganization of the Cytoskeleton},
  journal = {J. Neurosci.},
  year = {2004},
  volume = {24},
  pages = {3002--3012},
  number = {12},
  doi = {10.1523/JNEUROSCI.4963-03.2004},
}


@ARTICLE{Xu2010,
  author = {B Xu  and J S Goldman and  V V Rymar and  C Forget and  P S Lo and S J Bull and E Vereker and  P A Barker and L E Trudeau and  A F Sadikot and  T E Kennedy},
  title = {Critical roles for the netrin receptor deleted in colorectal cancer in dopaminergic neuronal precursor migration, axon guidance, and axon arborization},
  journal = {Neuroscience},
  year = {2004},
  volume = {169},
  pages = {932--49},
  number = {2},
  doi = {10.1016/j.neuroscience.2010.05.025},
}

@ARTICLE{Dun2017,
  author = {X. P. Dun and  D. B. Parkinson},
  title = {Role of Netrin-1 Signaling in Nerve Regeneration},
  journal = {Int. J. Mol. Sci.},
  year = {2017},
  volume = {18},
  pages = {491},
  number = {2},
  doi = {10.3390/ijms18030491},
}

@ARTICLE{Hinck2004,
  author = {L. Hinck},
  title = {The versatile roles of "axon guidance" cues in tissue morphogenesis},
  journal = {Dev. Cell},
  year = {2004},
  volume = {7},
  pages = {783-93},
  number = {6},
  doi = {10.1016/j.devcel.2004.11.002},
}


@ARTICLE{Baker2006,
  author = {K. A. Baker and S. W. Moore and A. A. Jarjour and T. E. Kennedy},
  title = {When a diffusible axon guidance cue stops diffusing: roles for netrins in adhesion and morphogenesis},
  journal = {Curr. Opin. Neurobiol.},
  year = {2006},
  volume = {16},
  pages = {529-34},
  number = {5},
  doi = {10.1016/j.conb.2006.08.002},
}


@ARTICLE{Hebrok2004,
author = {Hebrok, Matthias and Reichardt, Louis F},
journal = {Trends Cell Biol.},
month = {apr},
number = {4},
pages = {153--5},
title = {{Brain meets pancreas: netrin, an axon guidance molecule, controls epithelial cell migration.}},
volume = {14},
year = {2004}
}

@article{Yebra2003,
author = {Yebra, Mayra and Montgomery, Anthony M.P. and Diaferia, Giuseppe R. and Kaido, Thomas and Silletti, Steve and Perez, Brandon and Just, Margaret L. and Hildbrand, Simone and Hurford, Rosemary and Florkiewicz, Elin and Tessier-Lavigne, Marc and Cirulli, Vincenzo},
doi = {10.1016/S1534-5807(03)00330-7},
issn = {15345807},
journal = {Dev. Cell},
number = {5},
pages = {695--707},
title = {{Recognition of the neural chemoattractant netrin-1 by integrins $\alpha$6$\beta$4 and $\alpha$3$\beta$1 regulates epithelial cell adhesion and migration}},
volume = {5},
year = {2003}
}

@article{Marcos2009,
author = {Marcos, S{\'{e}}verine and Backer, St{\'{e}}phanie and Causeret, Fr{\'{e}}d{\'{e}}ric and Tessier-Lavigne, Marc and Bloch-Gallego, Evelyne},
doi = {10.1016/j.mcn.2009.04.008},
issn = {1095-9327 (Electronic)},
journal = {Mol. Cell. Neurosci.},
keywords = {Animals,Axons,Cell Movement,DCC Receptor,Embryo, Mammalian,Gene Expression Regulation, Developmental,Mice,Mice, Transgenic,Mutation,Nerve Growth Factors,Nerve Tissue Proteins,Netrin-1,Neural Pathways,Neurogenesis,Neurons,Olivary Nucleus,Organ Culture Techniques,Receptors, Cell Surface,Tumor Suppressor Proteins,cytology,embryology,genetics,metabolism,physiology},
language = {eng},
month = {aug},
number = {4},
pages = {429--439},
title = {{Differential roles of Netrin-1 and its receptor DCC in inferior olivary neuron  migration.}},
volume = {41},
year = {2009}
}

@article{Rappaz2016,
author = {Rappaz, Benjamin and {Lai Wing Sun}, Karen and Correia, James P and Wiseman, Paul W and Kennedy, Timothy E},
doi = {10.1371/journal.pone.0159405},
issn = {1932-6203 (Electronic)},
journal = {PloS one},
keywords = {Animals,Cells, Cultured,DCC Receptor,Fluorescence Resonance Energy Transfer,Growth Cones,Microdissection,Microscopy, Fluorescence,Nerve Growth Factors,Netrin-1,Rats, Sprague-Dawley,Receptors, Cell Surface,Signal Transduction,Spinal Cord,Tumor Suppressor Proteins,analysis,cdc42 GTP-Binding Protein,cytology,embryology,metabolism,methods,src-Family Kinases,ultrastructure},
language = {eng},
number = {8},
pages = {e0159405},
title = {{FLIM FRET Visualization of Cdc42 Activation by Netrin-1 in Embryonic Spinal  Commissural Neuron Growth Cones.}},
volume = {11},
year = {2016}
}

@article{Chen2017,
author = {Chen, Jing Ying and He, Xiao Xiao and Ma, Chi and Wu, Xin Min and Wan, Xi Lin and Xing, Zhen Kai and Pei, Qing Qing and Dong, Xian Ping and Liu, Dong Xu and Xiong, Wen Cheng and Zhu, Xiao Juan},
doi = {10.1038/s41598-017-05707-0},
issn = {20452322},
journal = {Sci. Rep.},
number = {1},
title = {{Netrin-1 promotes glioma growth by activating NF-$\kappa$B via UNC5A}},
volume = {7},
year = {2017}
}

@article{Fitamant2008,
author = {Fitamant, Julien and Guenebeaud, C{\'{e}}line and Coissieux, Marie May and Guix, Catherine and Treilleux, Isabelle and Scoazec, Jean Yves and Bachelot, Thomas and Bernet, Agn{\`{e}}s and Mehlen, Patrick},
doi = {10.1073/pnas.0709810105},
issn = {00278424},
journal = {Proc. Natl. Acad. Sci. U.S.A.},
keywords = {Apoptosis,DCC,Dependence receptor},
number = {12},
pages = {4850--4855},
title = {{Netrin-1 expression confers a selective advantage for tumor cell survival in metastatic breast cancer}},
volume = {105},
year = {2008}
}

@article{Ylivinkka2016,
title = {Netrin-1: A regulator of cancer cell motility?},
journal = {European Journal of Cell Biology},
volume = {95},
number = {11},
pages = {513-520},
year = {2016},
note = {Integrated mechano-chemical signals in invasion},
issn = {0171-9335},
doi = {10.1016/j.ejcb.2016.10.002},
author = {Irene Ylivinkka and Jorma Keski-Oja and Marko Hyyti{\"{a}}inen}
}


@article{Yamagishi2015,
author = {Yamagishi, Satoru and Yamada, Kohei and Sawada, Masato and Nakano, Suguru and Mori, Norio and Sawamoto, Kazunobu and Sato, Kohji},
doi = {10.3389/fncel.2015.00146},
issn = {16625102},
journal = {Front. Cell. Neurosci.},
keywords = {Adult neurogenesis,Axon guidance,Netrin,Rostral migratory stream,Subgranular zone,Subventricular zone},
number = {APR},
title = {{Netrin-5 is highly expressed in neurogenic regions of the adult brain}},
volume = {9},
year = {2015}
}

@article{Banyai1999,
author = {B{\'{a}}nyai, L{\'{a}}szl{\'{o}} and Patthy, L{\'{a}}szl{\'{o}}},
doi = {10.1110/ps.8.8.1636},
issn = {0961-8368},
journal = {Protein Sci.},
number = {8},
pages = {1636--1642},
title = {{The NTR module: Domains of netrins, secreted frizzled related proteins, and type I procollagen C‐proteinase enhancer protein are homologous with tissue inhibitors of metalloproteases}},
volume = {8},
year = {1999}
}

@misc{DeWit2007,
author = {{De Wit}, Joris and Verhaagen, Joost},
booktitle = {Advances in Experimental Medicine and Biology},
doi = {10.1007/978-0-387-70956-7_7},
isbn = {9780387709550},
issn = {00652598},
pages = {73--89},
title = {{Proteoglycans as modulators of axon guidance cue function}},
volume = {600},
year = {2007},
editor = {Gerard Pasterkamp},
publisher = {Springer, New York, N.Y}
}

@article{Fearon1990,
author = {Fearon, Eric R. and Vogelstein, Bert},
title = {A genetic model for colorectal tumorigenesis},
journal = {Cell},
volume = {61},
number = {5},
pages = {759-767},
year = {1990},
issn = {0092-8674},
doi = {10.1016/0092-8674(90)90186-I}
}


@article{Hedrick1994,
author = {Hedrick, L and Cho, K R and Fearon, E R and Wu, T C and Kinzler, K W and Vogelstein, B},
doi = {10.1101/gad.8.10.1174},
issn = {0890-9369 (Print)},
journal = {Genes Dev.},
keywords = {Adenocarcinoma, Mucinous,Adenoma,Adenomatous Polyps,Animals,Carcinoma,Cell Adhesion Molecules,Cell Differentiation,Cell Transformation, Neoplastic,Colorectal Neoplasms,DCC Receptor,DNA, Complementary,Gene Expression Regulation, Neoplastic,Genes, DCC,Humans,Intestinal Polyps,Molecular Sequence Data,Open Reading Frames,Organ Specificity,RNA, Messenger,Rats,Receptors, Cell Surface,Recombinant Fusion Proteins,Tumor Cells, Cultured,Tumor Suppressor Proteins,analysis,biosynthesis,genetics,metabolism,pathology},
language = {eng},
month = {may},
number = {10},
pages = {1174--1183},
title = {{The DCC gene product in cellular differentiation and colorectal tumorigenesis.}},
volume = {8},
year = {1994}
}

@article{Mehlen1998,
author = {Mehlen, Patrick and Rabizadeh, Shahrooz and Snipas, Scott J. and Assa-Munt, Nuria and Salvesen, Guy S. and Bredesen, Dale E.},
doi = {10.1038/27441},
issn = {00280836},
journal = {Nature},
number = {6704},
pages = {801--804},
title = {{The DCC gene product induces apoptosis by a mechanism requiring receptor proteolysis}},
volume = {395},
year = {1998}
}

@article{Lim2002,
author = {Lim, Yoo-shick and Wadsworth, William G},
doi = {10.1523/JNEUROSCI.22-16-07080.2002},
issn = {1529-2401 (Electronic)},
journal = {J. Neurosci.},
keywords = {Alleles,Amino Acid Motifs,Animals,Animals, Genetically Modified,Axons,Caenorhabditis elegans,Caenorhabditis elegans Proteins,Cell Movement,Growth Cones,Helminth Proteins,Mutagenesis, Site-Directed,Nerve Growth Factors,Nerve Tissue Proteins,Netrins,Phenotype,Protein Structure, Tertiary,Sequence Deletion,Structure-Activity Relationship,Transgenes,chemistry,genetics,physiology,ultrastructure},
language = {eng},
month = {aug},
number = {16},
pages = {7080--7087},
title = {{Identification of domains of netrin UNC-6 that mediate attractive and repulsive  guidance and responses from cells and growth cones.}},
volume = {22},
year = {2002}
}

@article{Xu2014,
author = {Xu, Kai and Wu, Zhuhao and Renier, Nicolas and Antipenko, Alexander and Tzvetkova-Robev, Dorothea and Xu, Yan and Minchenko, Maria and Nardi-Dei, Vincenzo and Rajashankar, Kanagalaghatta R. and Himanen, Juha and Tessier-Lavigne, Marc and Nikolov, Dimitar B.},
doi = {10.1126/science.1255149},
issn = {10959203},
journal = {Science},
number = {6189},
pages = {1275--1279},
title = {{Structures of netrin-1 bound to two receptors provide insight into its axon guidance mechanism}},
volume = {344},
year = {2014}
}

@article{Finci2014,
author = {Finci, Lorenzo I. and Kr{\"{u}}ger, Nina and Sun, Xiaqin and Zhang, Jie and Chegkazi, Magda and Wu, Yu and Schenk, Gundolf and Mertens, Haydyn D.T. and Svergun, Dmitri I. and Zhang, Yan and huai Wang, Jia and Meijers, Rob},
doi = {10.1016/j.neuron.2014.07.010},
issn = {10974199},
journal = {Neuron},
number = {4},
pages = {839--849},
pmid = {25123307},
title = {{The Crystal Structure of Netrin-1 in Complex with DCC Reveals the Bifunctionality of Netrin-1 As a Guidance Cue}},
volume = {83},
year = {2014}
}


@article{Geisbrecht2003,
author = {Geisbrecht, Brian V and Dowd, Kimberly A and Barfield, Ronald W and Longo, Patti A and Leahy, Daniel J},
doi = {10.1074/jbc.M302943200},
issn = {0021-9258 (Print)},
journal = {J. Biol. Chem.},
keywords = {Algorithms,Animals,Binding Sites,CHO Cells,Cell Adhesion Molecules,Cell Line,Chromatography, Gel,Cricetinae,Cross-Linking Reagents,DCC Receptor,Dose-Response Relationship, Drug,Electrophoresis, Polyacrylamide Gel,Enzyme-Linked Immunosorbent Assay,Heparin,Humans,Immunoblotting,Microscopy, Fluorescence,Models, Biological,Mutagenesis, Site-Directed,Nerve Growth Factors,Netrin Receptors,Netrin-1,Plasmids,Polymerase Chain Reaction,Protein Binding,Protein Structure, Tertiary,Receptors, Cell Surface,Transfection,Tumor Suppressor Proteins,chemistry,metabolism,pharmacology},
language = {eng},
month = {aug},
number = {35},
pages = {32561--32568},
title = {{Netrin binds discrete subdomains of DCC and UNC5 and mediates interactions  between DCC and heparin.}},
volume = {278},
year = {2003}
}

@article{Finci2015,
author = {Finci, L and Zhang, Y and Meijers, R and Wang, J-H},
doi = {10.1016/j.pbiomolbio.2015.04.001},
issn = {1873-1732 (Electronic)},
journal = {Prog. Biophys. Mol. Biol.},
keywords = {Animals,Axons,Cytoplasm,DCC Receptor,Humans,Protein Structure, Tertiary,Receptors, Cell Surface,Signal Transduction,Tumor Suppressor Proteins,chemistry,metabolism},
language = {eng},
month = {sep},
number = {3},
pages = {153--160},
title = {{Signaling mechanism of the netrin-1 receptor DCC in axon guidance.}},
volume = {118},
year = {2015}
}


@article{Gopal2016,
author = {Gopal, Angelica A and Rappaz, Benjamin and Rouger, Vincent and Martyn, Iain B and Dahlberg, Peter D and Meland, Rachel J and Beamish, Ian V and Kennedy, Timothy E and Wiseman, Paul W},
doi = {10.1016/j.bpj.2015.12.022},
issn = {1542-0086 (Electronic)},
journal = {Biophys. J.},
keywords = {Cell Membrane,DCC Receptor,HEK293 Cells,Humans,Nerve Growth Factors,Netrin Receptors,Netrin-1,Protein Transport,Receptors, Cell Surface,Tumor Suppressor Proteins,metabolism},
language = {eng},
month = {feb},
number = {3},
pages = {623--634},
title = {{Netrin-1-Regulated Distribution of UNC5B and DCC in Live Cells Revealed by TICCS.}},
volume = {110},
year = {2016}
}

@article{Bouchard2004,
author = {Bouchard, Jean Fran{\c{c}}ois and Moore, Simon W. and Tritsch, Nicolas X. and Roux, Philippe P. and Shekarabi, Masoud and Barker, Philip A. and Kennedy, Timothy E.},
doi = {10.1523/JNEUROSCI.4934-03.2004},
issn = {02706474},
journal = {J. Neurosci.},
keywords = {Endocytosis,Exocytosis,Growth cone,Outgrowth,Regeneration,cAMP},
month = {mar},
number = {12},
pages = {3040--3050},
title = {{Protein Kinase A Activation Promotes Plasma Membrane Insertion of DCC from An Intracellular Pool: A Novel Mechanism Regulating Commissural Axon Extension}},
volume = {24},
year = {2004}
}

@article{Matsumoto2010,
author = {Matsumoto, Hideko and Nagashima, Masabumi},
doi = {10.1016/j.neures.2010.02.004},
issn = {01680102},
journal = {Neurosci. Res.},
keywords = {Axon guidance,Axon shaft,Cortical development,DCC,Exocytosis,Netrin-1},
month = {jun},
number = {2},
pages = {99--107},
title = {{Netrin-1 elevates the level and induces cluster formation of its receptor DCC at the surface of cortical axon shafts in an exocytosis-dependent manner}},
volume = {67},
year = {2010}
}

@article{Forcet2001,
author = {Forcet, C. and Ye, X. and Granger, L. and Corset, V. and Shin, H. and Bredesen, D. E. and Mehlen, P.},
doi = {10.1073/pnas.051378298},
issn = {0027-8424},
journal = {Proc. Natl. Acad. Sci. U.S.A.},
title = {{The dependence receptor DCC (deleted in colorectal cancer) defines an alternative mechanism for caspase activation}},
year = {2001}
}

@article{Ly2008,
author = {Ly, Alice and Nikolaev, Anatoly and Suresh, Geetha and Zheng, Yufang and Tessier-Lavigne, Marc and Stein, Elke},
doi = {10.1016/j.cell.2008.05.030},
file = {::},
issn = {1097-4172},
journal = {Cell},
keywords = {Animals,Axons,Axons: metabolism,COS Cells,Cercopithecus aethiops,Embryo, Mammalian,Embryo, Mammalian: metabolism,Membrane Proteins,Membrane Proteins: chemistry,Membrane Proteins: metabolism,Nerve Growth Factors,Nerve Growth Factors: metabolism,Protein Structure, Tertiary,Rats,Receptors, Cell Surface,Receptors, Cell Surface: metabolism,Spinal Cord,Spinal Cord: cytology,Spinal Cord: embryology,Spinal Cord: metabolism,Tumor Suppressor Proteins,Tumor Suppressor Proteins: metabolism,Xenopus,ly:2008},
mendeley-tags = {ly:2008},
month = {jun},
number = {7},
pages = {1241--54},
title = {{DSCAM is a netrin receptor that collaborates with DCC in mediating turning responses to netrin-1.}},
volume = {133},
year = {2008}
}

@article{Guirland2004,
author = {Guirland, Carmine and Suzuki, Shingo and Kojima, Masami and Lu, Bai and Zheng, James Q.},
doi = {10.1016/S0896-6273(04)00157-6},
issn = {08966273},
journal = {Neuron},
month = {apr},
number = {1},
pages = {51--62},
title = {{Lipid rafts mediate chemotropic guidance of nerve growth cones}},
volume = {42},
year = {2004}
}

@article{Herincs2005,
author = {H{\'{e}}rincs, Zolt{\'{a}}n and Corset, V{\'{e}}ronique and Cahuzac, Nathalie and Furne, C{\'{e}}line and Castellani, Val{\'{e}}rie and Hueber, Anne-Odile and Mehlen, Patrick},
doi = {10.1242/jcs.02296},
issn = {0021-9533},
journal = {J. Cell Sci.},
keywords = {Animals,Cell Adhesion Molecules,Cell Adhesion Molecules: metabolism,Cell Communication,Cell Communication: physiology,Cell Line,Cells, Cultured,Chemotaxis,Chemotaxis: physiology,Cues,Enzyme Activation,Enzyme Activation: physiology,Extracellular Signal-Regulated MAP Kinases,Extracellular Signal-Regulated MAP Kinases: metabo,Growth Cones,Growth Cones: metabolism,Humans,Membrane Lipids,Membrane Lipids: metabolism,Membrane Microdomains,Membrane Microdomains: metabolism,Nerve Growth Factors,Nerve Growth Factors: metabolism,Palmitic Acid,Palmitic Acid: metabolism,Rats,Receptors, Cell Surface,Signal Transduction,Signal Transduction: physiology,Spinal Cord,Spinal Cord: embryology,Tumor Suppressor Proteins,Tumor Suppressor Proteins: metabolism,herincs:2005},
mendeley-tags = {herincs:2005},
month = {apr},
number = {Pt 8},
pages = {1687--92},
title = {{DCC association with lipid rafts is required for netrin-1-mediated axon guidance.}},
volume = {118},
year = {2005}
}


@article{Petrie2009,
author = {Petrie, Ryan J and Zhao, Beibei and Bedford, Fiona and Lamarche-Vane, Nathalie},
doi = {10.1042/BC20070108},
issn = {1768-322X},
journal = {Biol. Cell},
keywords = {Animals,Brain,Brain: metabolism,Cell Line, Tumor,Humans,Membrane Microdomains,Membrane Microdomains: genetics,Membrane Microdomains: metabolism,Nerve Growth Factors,Nerve Growth Factors: genetics,Nerve Growth Factors: metabolism,Neuroblastoma,Neuroblastoma: genetics,Neuroblastoma: metabolism,Protein Binding,Protein Transport,Rats,Receptors, Cell Surface,Receptors, Cell Surface: genetics,Receptors, Cell Surface: metabolism,Signal Transduction,Tumor Suppressor Proteins,Tumor Suppressor Proteins: genetics,Tumor Suppressor Proteins: metabolism,petrie:2009},
mendeley-tags = {petrie:2009},
month = {feb},
number = {2},
pages = {77--90},
title = {{Compartmentalized DCC signalling is distinct from DCC localized to lipid rafts.}},
volume = {101},
year = {2009}
}

@article{Chang2007,
author = {Chang, Veronica T and Crispin, Max and Aricescu, A Radu and Harvey, David J and Nettleship, Joanne E and Fennelly, Janet A and Yu, Chao and Boles, Kent S and Evans, Edward J and Stuart, David I and Dwek, Raymond A and Jones, E Yvonne and Owens, Raymond J and Davis, Simon J},
doi = {10.1016/j.str.2007.01.011},
issn = {0969-2126 (Print)},
journal = {Structure (London, England : 1993)},
keywords = {Alkaloids,Cell Line,Cloning, Molecular,Enzyme Inhibitors,Genomics,Glycoproteins,Glycosylation,Humans,Swainsonine,antagonists & inhibitors,chemistry,drug effects,genetics,metabolism,methods},
language = {eng},
month = {mar},
number = {3},
pages = {267--273},
title = {{Glycoprotein structural genomics: solving the glycosylation problem.}},
volume = {15},
year = {2007}
}


@article{Grandin2016,
author = {Grandin, M{\'{e}}lodie and Meier, Markus and Delcros, Jean Guy and Nikodemus, Denise and Reuten, Raphael and Patel, Trushar R. and Goldschneider, David and Orriss, George and Krahn, Natalie and Boussouar, Amina and Abes, Riad and Dean, Yann and Neves, David and Bernet, Agnes and Depil, Stephane and Schneiders, Fiona and Poole, Kate and Dante, Robert and Koch, Manuel and Mehlen, Patrick and Stetefeld, J{\"{o}}rg},
doi = {10.1016/j.ccell.2016.01.001},
issn = {18783686},
journal = {Cancer Cell},
month = {feb},
number = {2},
pages = {173--185},
publisher = {Cell Press},
title = {{Structural Decoding of the Netrin-1/UNC5 Interaction and its Therapeutical Implications in Cancers}},
volume = {29},
year = {2016}
}


@phdthesis{Kruger2014,
  title={Structural and functional analysis of the guidance cue molecule netrin-1 in complex with its receptor DCC unravels the molecular mechanisms of interaction},
  author={Kr{\"u}ger, Nina},
  year={2014},
  school={Staats-und Universit{\"a}tsbibliothek Hamburg Carl von Ossietzky}
}


@article{Serafini1994,
author = {Serafini, T and Kennedy, T E and Galko, M J and Mirzayan, C and Jessell, T M and Tessier-Lavigne, M},
doi = {10.1016/0092-8674(94)90420-0},
issn = {0092-8674 (Print)},
journal = {Cell},
keywords = {Amino Acid Sequence,Animals,Axons,Base Sequence,Brain Chemistry,Caenorhabditis elegans Proteins,Chick Embryo,Cloning, Molecular,Dose-Response Relationship, Drug,Drug Synergism,Helminth Proteins,Molecular Sequence Data,Nerve Growth Factors,Nerve Tissue Proteins,Netrin-1,Netrins,Peptide Fragments,Recombinant Proteins,Sequence Analysis,Sequence Homology, Amino Acid,Spinal Cord,Tumor Suppressor Proteins,chemistry,cytology,drug effects,genetics,growth & development,isolation & purification,pharmacology,physiology},
language = {eng},
month = {aug},
number = {3},
pages = {409--424},
title = {{The netrins define a family of axon outgrowth-promoting proteins homologous to C.  elegans UNC-6.}},
volume = {78},
year = {1994}
}

@article{Kappler2000,
author = {Kappler, J and Franken, S and Junghans, U and Hoffmann, R and Linke, T and M\"uller, H W and Koch, K W},
doi = {10.1006/bbrc.2000.2583},
issn = {0006-291X (Print)},
journal = {Biochem. Biophys. Res. Commun.},
keywords = {Amino Acid Sequence,Animals,Biotinylation,Calorimetry,Circular Dichroism,Glycosaminoglycans,Heparin,Kinetics,Molecular Sequence Data,Nerve Growth Factors,Netrin-1,Peptides,Protein Binding,Protein Structure, Secondary,Surface Plasmon Resonance,Temperature,Time Factors,Tumor Suppressor Proteins,chemistry,metabolism},
language = {eng},
month = {may},
number = {2},
pages = {287--291},
title = {{Glycosaminoglycan-binding properties and secondary structure of the C-terminus of  netrin-1.}},
volume = {271},
year = {2000}
}


@article{Tessier-Lavigne1996,
author = {Tessier-Lavigne, M and Goodman, C S},
doi = {10.1126/science.274.5290.1123},
issn = {0036-8075 (Print)},
journal = {Science (New York, N.Y.)},
keywords = {Animals,Axons,Biological Evolution,Cell Adhesion Molecules, Neuronal,Cell Communication,Cell Movement,Humans,Ligands,Nerve Growth Factors,Nerve Tissue Proteins,Nervous System,Neural Pathways,Receptors, Cell Surface,embryology,physiology},
language = {eng},
month = {nov},
number = {5290},
pages = {1123--1133},
title = {{The molecular biology of axon guidance.}},
volume = {274},
year = {1996}
}

@article{Stanco2009,
author = {Stanco, Amelia and Szekeres, Charles and Patel, Nikhil and Rao, Sarada and Campbell, Kenneth and Kreidberg, Jordan A and Polleux, Franck and Anton, E S},
doi = {10.1073/pnas.0811343106},
file = {::},
issn = {1091-6490},
journal = {Proc. Natl. Acad. Sci. U.S.A.},
keywords = {Animals,Cell Movement,Cell Movement: genetics,Cerebral Cortex,Cerebral Cortex: cytology,Cerebral Cortex: growth & development,Cerebral Cortex: metabolism,Integrin alpha3beta1,Integrin alpha3beta1: metabolism,Interneurons,Interneurons: metabolism,Interneurons: physiology,Mice,Mice, Knockout,Nerve Growth Factors,Nerve Growth Factors: genetics,Nerve Growth Factors: metabolism,Tumor Suppressor Proteins,Tumor Suppressor Proteins: genetics,Tumor Suppressor Proteins: metabolism,gamma-Aminobutyric Acid,gamma-Aminobutyric Acid: metabolism,stanco:2009},
mendeley-tags = {stanco:2009},
month = {may},
number = {18},
pages = {7595--600},
title = {{Netrin-1-alpha3beta1 integrin interactions regulate the migration of interneurons through the cortical marginal zone.}},
volume = {106},
year = {2009}
}


@article{Nikolopoulos2005,
author = {Nikolopoulos, Sotiris N and Giancotti, Filippo G},
issn = {1551-4005},
journal = {Cell Cycle},
keywords = {Animals,Axons,Axons: metabolism,Binding Sites,Body Patterning,Cell Differentiation,Cell Movement,Epithelium,Epithelium: metabolism,Focal Adhesion Protein-Tyrosine Kinases,Focal Adhesion Protein-Tyrosine Kinases: metabolis,Humans,Integrins,Integrins: metabolism,Ligands,Mice,Models, Biological,Neovascularization, Physiologic,Nerve Growth Factors,Nerve Growth Factors: metabolism,Neurons,Neurons: metabolism,Protein Binding,Proto-Oncogene Proteins c-fyn,Proto-Oncogene Proteins c-fyn: metabolism,Receptors, Cell Surface,Receptors, Cell Surface: metabolism,Signal Transduction,Tumor Suppressor Proteins,Tumor Suppressor Proteins: metabolism,nikolopoulos:2005},
mendeley-tags = {nikolopoulos:2005},
month = {mar},
number = {3},
pages = {e131--5},
title = {{Netrin-integrin signaling in epithelial morphogenesis, axon guidance and vascular patterning.}},
volume = {4},
year = {2005}
}

@article{Moore2009,
author = {Moore, Simon W and Biais, Nicolas and Sheetz, Michael P},
doi = {10.1126/science.1173851},
file = {::},
issn = {1095-9203},
journal = {Science},
keywords = {Animals,Axons,Axons: physiology,Biomechanics,Cells, Cultured,Cues,Growth Cones,Growth Cones: physiology,Immobilized Proteins,Immobilized Proteins: physiology,Mice,Nerve Growth Factors,Nerve Growth Factors: physiology,Neurons,Neurons: physiology,Receptors, Cell Surface,Receptors, Cell Surface: physiology,Spinal Cord,Spinal Cord: cytology,Spinal Cord: embryology,Tumor Suppressor Proteins,Tumor Suppressor Proteins: physiology,moore:2009},
mendeley-tags = {moore:2009},
month = {jul},
number = {5937},
pages = {166},
title = {{Traction on immobilized netrin-1 is sufficient to reorient axons.}},
volume = {325},
year = {2009}
}

@article{Kennedy1994,
author = {Kennedy, Timothy E. and Serafini, Tito and de la Torre, Jos{\'{e}}R R. and Tessier-Lavigne, Marc},
doi = {10.1016/0092-8674(94)90421-9},
issn = {00928674},
journal = {Cell},
title = {{Netrins are diffusible chemotropic factors for commissural axons in the embryonic spinal cord}},
year = {1994}
}

@article{Causeret2002,
author = {Causeret, Fr{\'{e}}d{\'{e}}ric and Danne, Fran{\c{c}}ois and Ezan, Fr{\'{e}}d{\'{e}}ric and Sotelo, Constantino and Bloch-Gallego, Evelyne},
doi = {10.1006/dbio.2002.0681},
issn = {00121606},
journal = {Dev. Biol.},
keywords = {Axon outgrowth,Brainstem,Chemotropic molecules,Collagen assays,Floor-plate,In situ hybridization,Mice,Nuclear translocation,Precerebellar nuclei,Robo receptors},
month = {jun},
number = {2},
pages = {429--440},
publisher = {Academic Press},
title = {{Slit antagonizes netrin-1 attractive effects during the migration of inferior olivary neurons}},
volume = {246},
year = {2002}
}

@article{Sprague2004,
author = {Sprague, Brian L. and Pego, Robert L. and Stavreva, Diana A. and McNally, James G.},
doi = {10.1529/biophysj.103.026765},
issn = {00063495},
journal = {Biophys. J.},
number = {6},
pages = {3473--3495},
title = {{Analysis of binding reactions by fluorescence recovery after photobleaching}},
volume = {86},
year = {2004}
}

@article{Serge2008,
author = {Serg{\'{e}}, Arnauld and Bertaux, Nicolas and Rigneault, Herv{\'{e}} and Marguet, Didier},
doi = {10.1038/nmeth.1233},
issn = {15487091},
journal = {Nat. Methods},
title = {{Dynamic multiple-target tracing to probe spatiotemporal cartography of cell membranes}},
year = {2008}
}

@article{Michalet2010,
author = {Michalet, Xavier},
doi = {10.1103/PhysRevE.82.041914},
issn = {1550-2376 (Electronic)},
journal = {Phys. Rev. E.},
keywords = {Models, Theoretical,Motion,Uncertainty},
language = {eng},
month = {oct},
number = {4 Pt 1},
pages = {41914},
title = {{Mean square displacement analysis of single-particle trajectories with  localization error: Brownian motion in an isotropic medium.}},
volume = {82},
year = {2010}
}

@article{Normanno2015,
author = {Normanno, Davide and Boudar{\`{e}}ne, Lydia and Dugast-Darzacq, Claire and Chen, Jiji and Richter, Christian and Proux, Florence and B{\'{e}}nichou, Olivier and Voituriez, Rapha{\"{e}}l and Darzacq, Xavier and Dahan, Maxime},
doi = {10.1038/ncomms8357},
issn = {2041-1723 (Electronic)},
journal = {Nat. Commun.},
keywords = {Base Sequence,Cell Line, Tumor,DNA-Binding Proteins,Gene Expression Regulation,Humans,Kinetics,Protein Conformation,Protein Transport,Repressor Proteins,genetics,metabolism,physiology},
language = {eng},
month = {jul},
pages = {7357},
title = {{Probing the target search of DNA-binding proteins in mammalian cells using TetR  as model searcher.}},
volume = {6},
year = {2015}
}

@article{Bohrer2021,
author = {Bohrer, Christopher H and Yang, Xinxing and Thakur, Shreyasi and Weng, Xiaoli and Tenner, Brian and McQuillen, Ryan and Ross, Brian and Wooten, Matthew and Chen, Xin and Zhang, Jin and Roberts, Elijah and Lakadamyali, Melike and Xiao, Jie},
issn = {1548-7105 (Electronic)},
journal = {Nat. Methods},
keywords = {Algorithms,Artifacts,Fluorescent Dyes,Microscopy, Fluorescence,Single Molecule Imaging,Stochastic Processes,chemistry,methods},
language = {eng},
month = {jun},
number = {6},
pages = {669--677},
title = {{A pairwise distance distribution correction (DDC) algorithm to eliminate  blinking-caused artifacts in SMLM.}},
volume = {18},
year = {2021},
doi = {10.1038/s41592-021-01154-y}
}


@article{Suzuki2006,
author = {Suzuki, Ryota and Shimodaira, Hidetoshi},
doi = {10.1093/bioinformatics/btl117},
issn = {1367-4803 (Print)},
journal = {Bioinformatics (Oxford, England)},
keywords = {Algorithms,Cluster Analysis,Computational Biology,Gene Expression Profiling,Gene Expression Regulation, Neoplastic,Humans,Internet,Models, Statistical,Oligonucleotide Array Sequence Analysis,Probability,Programming Languages,Software,Uncertainty,methods},
language = {eng},
month = {jun},
number = {12},
pages = {1540--1542},
title = {{Pvclust: an R package for assessing the uncertainty in hierarchical clustering.}},
volume = {22},
year = {2006}
}

@article{Tran2013,
author = {Tran, Thanh N. and Drab, Klaudia and Daszykowski, Michal},
doi = {10.1016/j.chemolab.2012.11.006},
issn = {01697439},
journal = {Chemom. Intell. Lab. Syst.},
keywords = {Clustering,Density-based clustering},
month = {jan},
pages = {92--96},
publisher = {Elsevier},
title = {{Revised DBSCAN algorithm to cluster data with dense adjacent clusters}},
volume = {120},
year = {2013}
}
